# 7-Dehydrocholesterol-derived oxysterols cause neurogenic defects in Smith-Lemli-Opitz syndrome

**DOI:** 10.1101/2021.01.30.428955

**Authors:** Hideaki Tomita, Kelly M. Hines, Josi M. Herron, Amy Li, David W. Baggett, Libin Xu

## Abstract

Defective 3β-hydroxysterol-Δ^7^ -reductase (DHCR7) in the developmental disorder, Smith-Lemli-Opitz syndrome (SLOS), results in deficiency in cholesterol and accumulation of its precursor, 7-dehydrocholesterol (7-DHC). Here, we show that loss of *DHCR7* causes accumulation of 7-DHC-derived oxysterol metabolites, premature neurogenesis, and perturbation of neuronal localization in developing murine or human cortical neural precursors, both *in vitro* and *in vivo*. We found that a major oxysterol, 3β,5*α*-dihydroxycholest-7-en-6-one (DHCEO), mediates these effects by initiating crosstalk between glucocorticoid receptor (GR) and neurotrophin receptor kinase TrkB. Either loss of *DHCR7* or direct exposure to DHCEO causes hyperactivation of GR and TrkB and their downstream MEK-ERK-C/EBP signaling pathway in cortical neural precursors. Moreover, direct inhibition of GR activation with an antagonist or inhibition of DHCEO accumulation with antioxidants rescues the premature neurogenesis phenotype caused by the loss of *DHCR7*. These results suggest that GR could be a new therapeutic target against the neurological defects observed in SLOS.

## Introduction

Brain is rich in cholesterol, contributing to 25% of total cholesterol in human body, and nearly all cholesterol in the brain is synthesized locally (Dietschy and Turley, 2004). Therefore, dysregulation of cholesterol metabolism in CNS can potentially cause significant defects in CNS development and functions. Indeed, enzymatic deficiencies in cholesterol biosynthesis cause a number of inherited diseases with severe neurodevelopmental phenotypes (Porter and Herman, 2011).

Smith-Lemli Opitz syndrome (SLOS) is an autosomal recessive, neurological and developmental disorder characterized by multiple developmental defects, such as distinctive facial features, cleft palate, microcephaly and holoprosencephaly, as well as severe intellectual impairment and behavioral problems (Porter and Herman, 2011; Thurm et al., 2016). Notably, SLOS patients display high incidence (> 50%) of autism spectrum disorders (Bukelis et al., 2007; Sikora et al., 2006; Tierney et al., 2006). SLOS is caused by mutations in the 3β-hydroxysterol-Δ^7^ -reductase gene (*DHCR7*), which encodes the enzyme that converts 7-dehydrocholesterol (7-DHC) to cholesterol in the final step of the cholesterol biosynthesis pathway (Fitzky et al., 1998; Tint et al., 1994; Wassif et al., 1998). Defective DHCR7 resulting from the mutations leads to deficiency in cholesterol and accumulation of 7-DHC in tissues and fluids of affected individuals (Tint et al., 1994; Tint et al., 1995). 7-DHC was found to be highly reactive toward free radical oxidation, leading to the formation of its oxidative metabolites, *i.e*., oxysterols (Xu et al., 2009; Xu et al., 2010; Xu et al., 2013; Xu et al., 2011). 7-DHC-derived oxysterols can exert cytotoxicity in neuronal cells, induce gene expression changes, and increase formation of dendritic arborization from cortical neurons (Korade et al., 2010; Xu et al., 2012). Interestingly, increased dendrite and axon formation has also been observed in neurons isolated from *Dhcr7*-KO mouse brain (Jiang et al., 2010). These studies suggest that 7-DHC-derived oxysterols may be underlying the alterations in neuronal processes in SLOS.

Cholesterol, 7-DHC, and oxysterols derived from both have been found to play important roles in modulating signaling pathways in developing tissues and organs, such as Hedgehog (Hh) and Wnt signaling pathways (Byrne et al., 2016; Corcoran and Scott, 2006; Huang et al., 2018; Myers et al., 2017; Porter et al., 1996; Raleigh et al., 2018). Related to SLOS, ring-B oxysterols derived from 7-DHC oxidation inhibit Smo in the Hh pathway (Sever et al., 2016). On the other hand, cholesterol was recently found to selectively activate canonical Wnt signaling over non-canonical Wnt signaling (Sheng et al., 2014). However, in human induced pluripotent stem cells (hiPSCs) derived from SLOS patient fibroblasts, accumulation of 7-DHC was found to inhibit Wnt/β-catenin pathway, which contributes to the precocious neuronal specification in SLOS neural progenitors (Francis et al., 2016). Depending on the position of the oxidation, oxysterols have also been shown to bind and activate other signaling molecules and nuclear receptors, including estrogen receptors, liver X receptors, and glucocorticoid receptor (GR), and thus, play important roles in neurodevelopment and diseases (DuSell et al., 2008; Theofilopoulos et al., 2013; Voisin et al., 2017). However, it is unknown how alteration of sterol composition influences neural stem cell/progenitor behaviors during cortical development, and it remains elusive whether neural defects of SLOS is due to deficiency in cholesterol or accumulation of 7-DHC or its oxysterols.

Here we examine the effects of *DHCR7* mutations in developing neural precursors, focusing on the cerebral cortex. We demonstrate that 7-DHC-derived oxysterols start to accumulate at embryonic day 12.5 (E12.5) and continue to increase at E14.5 and E16.5, and that loss of *DHCR7* causes decreased proliferation and self-renewal of mouse cortical neural precursors and aberrant premature neurogenesis in both mouse and human neural progenitor cells (NPCs). We then provide evidence that a 7-DHC-derived oxysterol, 3β,5*α*-dihydroxycholest-7-en-6-one (DHCEO), activates GR and the downstream receptor tyrosine kinases (RTKs)-mediated neurogenic signaling through TrkB, and in doing so, promotes premature NPC differentiation and perturbs neuronal positioning. Either inhibition of GR activation with an antagonist or inhibition of DHCEO accumulation with antioxidants rescues the premature neurogenesis defect.

## Results

### Dhcr7 is Expressed in Embryonic Cortical Precursors and Neurons of Developing Murine Cortex

To understand the role of *Dhcr7* in neural development, we studied murine developing cortical precursors during embryonic neurogenesis. We first analyzed the expression of *Dhcr7* mRNA during mouse cortical development. RT-PCR analysis showed expression of *Dhcr7* in embryonic day 11 (E11) through postnatal day 0 (P0) cortex (Figure 1A). Quantitative PCR (qPCR) also suggested the stable expression of *Dhcr7* from E11 to E18 (Figure 1B). Western blot analysis further confirmed the presence of Dhcr7 protein in developing cortex (Figure 1C). Immunostaining showed that Dhcr7 was broadly expressed in the developing cortex and was detectable in precursors in the E12 ventricular and subventricular (VZ/SVZ) zones (Figure 1D). Immunostaining of cortical sections of *Dhcr7*-KO mouse brain at E12 with this particular Dhcr7 antibody confirmed the antibody specificity against Dhcr7. To further establish the expression of Dhcr7 in neural lineage cells, we analyzed cortical precursor cultures from E12.5, which consists of proliferating radial glial cells that generate neurons in culture (Figure 1E). In the E12.5 cortical precursor cultures, Dhcr7+ cells were co-labeled with the precursor markers, Sox2 and Pax6. Dhcr7 was also co-labeled with the neuronal marker, βIII-tubulin, and the neural lineage marker, Nestin. The results suggested that Dhcr7 was consistently expressed in neural lineage cells, which is consistent with our *in vivo* analysis.

**Figure 1.**
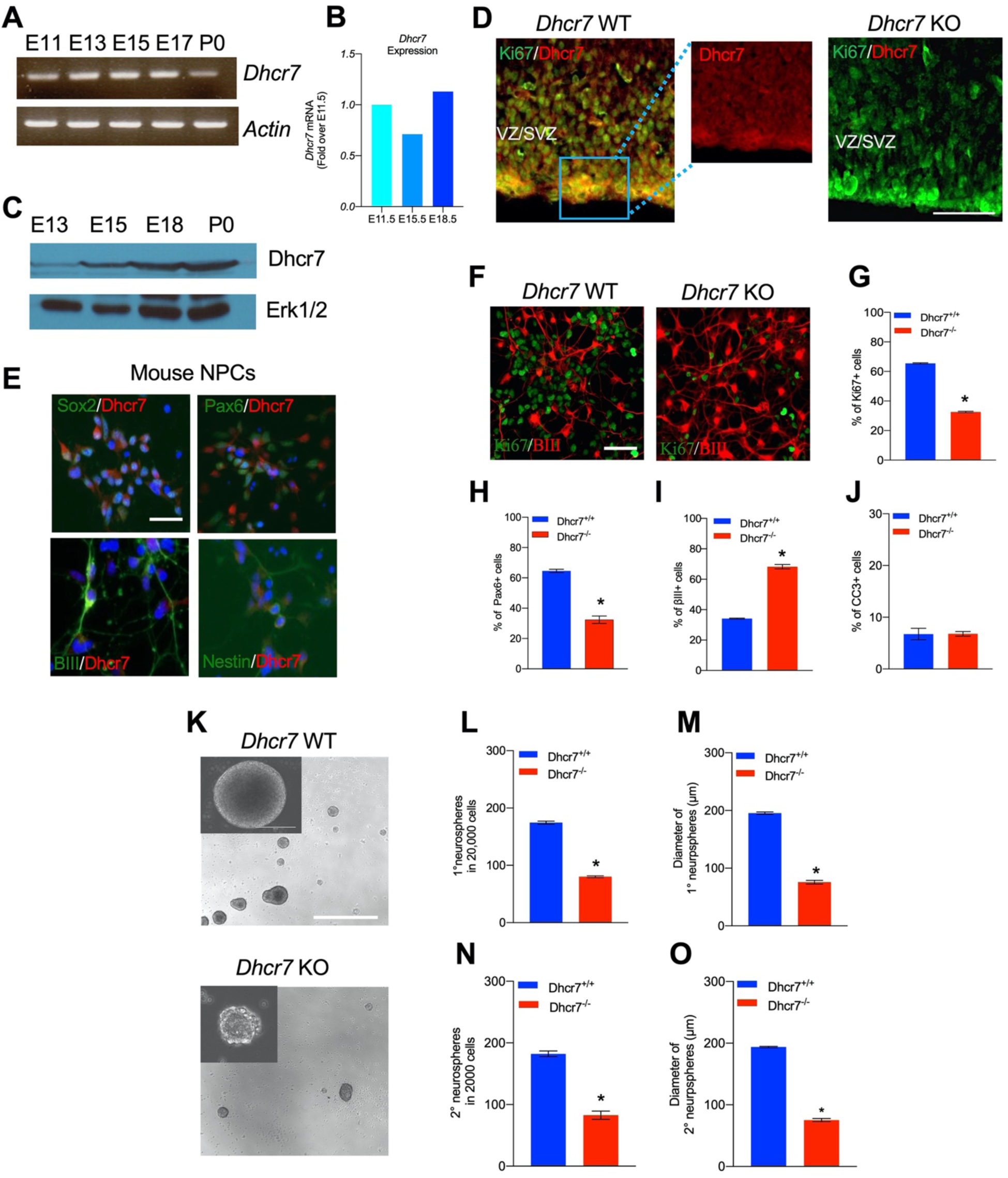
Loss of Dhcr7 alleles causes decreased proliferation and increased neurogenesis in murine cortical precursors. (A) RT-PCR for *Dhcr7* mRNA in the E11.5 to P0 cortex. *β-actin* mRNA was used as loading control. (B) qRT-PCR for *Dhcr7* mRNA in the E11.5 to E18.5 cortex. Data is expressed as fold change over E11.5 cortex. (C) Western blot of Dhcr7 in total cortical lysates from E13.5 to P0. The blot was re-probed for Erk1/2 as a loading control. (D) Images of E13.5 *Dhcr7^+/+^* (*Dhcr7*-WT, left panel) and *Dhcr7^-/-^* (*Dhcr7*-KO, right panel) cortical sections immunostained for Dhcr7 (red). The subventricular/ventricular zone (SVZ/VZ) is denoted. The right panels show image of boxed area. Scale bar =100 μm. (E) Images of cultured mouse cortical precursors immunostained for Dhcr7 (red) and Sox2, Pax6, βIII-tubulin and Nestin (green) and counterstained with DAPI (blue). Scale bar = 50 μm. (F-J) E12.5 cortical precursors from single *Dhcr^+/+^* and *Dhcr7^-/-^* embryos were cultured 3 days and analyzed. (F) Cells were immunostained for Ki67 (green) and βIII-tubulin (red) after 3 days and quantified for the proportions of Ki67+ (G), Pax6+ (H), βIII-tubulin+ (I) and CC3+ cells (J). Scale bar = 50 μm. *, *p* < 0.001; n = 3 embryos per genotype. (K-O) E13.5 cortical precursor cells from single *Dhcr7^+/+^* or *Dhcr7^-/-^* embryos were cultured as primary neurosphere (K) and the number and diameter of primary neurospheres were quantified 6 days later (L, M). Equal numbers of primary neurospheres were then passaged, and the number and diameter of secondary neurospheres were quantified 6 days later (N, O). Representative images of *Dhcr^+/+^* and *Dhcr7^-/-^* neurospheres are shown as inserts in the left corner. *, *p* < 0.001; n = 3 embryos per genotype. Scale Bar = 100 μm. **Figure 1-Figure Supplement 1.** Characterization of the pluripotency of SLOS-derived human iPSCs. **Figure 1-Figure Supplement 2.** Loss of *DHCR7* alleles causes decreased proliferation and increased neurogenesis in human cortical precursors.

### Loss of Dhcr7 Alleles Causes Decreased Proliferation and Increased Neurogenesis in Murine and Human Cortical Precursors

SLOS is characterized by neurodevelopmental defects, such as microcephaly. To ask if *Dhcr7* plays important roles in neural precursor (progenitor) development as seen in human patients with SLOS, we intercrossed *Dhcr7*^+/-^ mice and prepared single embryo cultures from E12.5 *Dhcr7*^-/-^ (knockout or KO) or *Dhcr7*^+/+^ embryos. Cultures were immunostained for Ki67 and βIII-tubulin 3 days after plating, which revealed that loss of *Dhcr7* caused significant decrease in the proportion of Ki67+ and Pax6+ precursors whereas increased the proportion of βIII-tubulin+ neurons (Figure 1F-I). Immunostaining for cleaved caspase 3 (CC3) showed that the loss of *Dhcr7* alleles did not affect the survival of cortical precursor cells in culture (Figure 1J). These results suggested that loss of *Dhcr7* leads to premature neurogenesis and decreased proliferation of cortical precursors.

To determine if Dhcr7 is important for the proliferation and self-renewal of cortical precursors, we performed neurosphere assays, which assess if sphere-forming precursors can self-renew and generate new spheres. E13.5 cortical precursors from *Dhcr7*^-/-^ and *Dhcr7*^+/+^ embryos were cultured in the presence of FGF2 and EGF. The number and diameter of spheres were measured 7 days post-plating (Figure 1K-O). Significantly fewer number of and smaller neurospheres were generated from *Dhcr7*^-/-^ embryonic cortices comparing to its wild-type littermates (Figure 1L-M). These results were consistent with the reduced proportion of Ki67+ precursors in the adherent cultures. Following formation of the primary neurospheres, these spheres were triturated and re-plated at equal density to form the secondary neurospheres. The results showed that there was an approximately 2-fold decrease in the number and diameter of secondary spheres from *Dhcr7*^-/-^ cortical precursors relative to *Dhcr7*^+/+^ (Figure 1N-O). Thus, the loss of *Dhcr7* alleles disrupts proliferation and self-renewal of cortical precursors.

We then asked whether *DHCR7* was also necessary for neurogenesis of human embryonic stem cell-derived NPCs. To examine function of DHCR7, we generated human induced pluripotent stem cells (hiPSCs) from two lines of SLOS patient fibroblasts and one line of wild-type fibroblasts and verified their pluripotency and stemness as described in Methods (Figure 1-Figure Supplement 1) (Okita et al., 2011; Yu et al., 2007). The SLOS hiPSCs were then differentiated into hNPCs as described previously. Immunoreactivity of DHCR7 was detected in almost all wild-type hNPCs in these cultures with Sox2+ and Nestin+ cells (Figure 1-Figure Supplement 2; Panel A). To examine neural differentiation, the SLOS and wild-type NPCs were differentiated in cultures and immunostained 4 days later for Ki67 and βIII-tubulin for proliferating precursors and newborn neurons, respectively (Figure 1-Figure Supplement 2; Panels B-D). Notably, the proportion of Ki67+ precursors was significantly decreased whereas the proportion of βIII-tubulin+ neurons was increased in SLOS NPC cultures, similar to the phenotype seen in murine cortical precursors from *Dhcr7*^-/-^ embryos. Taken together, DHCR7 is involved in proliferation and differentiation of human NPCs.

### Cholesterol precursor 7-DHC and 7-DHC-derived Oxysterols are accumulated in Dhcr7^-/-^ developing cortex and human NPCs

7-DHC is highly susceptible to free radical oxidation (Xu et al., 2009), which leads to formation of numerous oxysterols in cells and tissues (Xu et al., 2013; Xu et al., 2011) (Figure 2A). Liquid chromatography-tandem mass spectrometry (LC-MS/MS) was performed on cortices from *Dhcr7*^-/-^ and *Dhcr7*^+/+^ embryos during cortical development. These analyses revealed significant accumulation of 7-DHC and reduction of cholesterol in the embryonic cortices from *Dhcr7*^-/-^ embryos throughout cortical development from E12.5 to E16.5 (Figure 2B). Increased levels of 7-dehydrodesmosterol (7-DHD), the precursor to desmosterol via Dhcr7, and 8-dehydrocholesterol (8-DHC) and decreased level of desmosterol in *Dhcr7*^-/-^ cortices are also consistent with the loss of the function of Dhcr7. Furthermore, 7-DHC oxysterols, such as DHCEO, 4*α*-hydroxy-7-DHC (4*α*-OH-7DHC), and 4β-hydroxy-7-DHC (4β-OH-7DHC) (Xu et al., 2013; Xu et al., 2011) showed substantial accumulation in the *Dhcr7*^-/-^ cortices, reaching up to 4-8 ng/mg of tissue, throughout the development (Figure 2A and 2C). Note that 4 ng/mg of tissue would translate into approximately 10 μM assuming the density of the brain is 1 g/mL. On the other hand, cholesterol-derived oxysterols, such as 24-keto-Chol, 24,25-epoxy-Chol, 24- or 25-OH-Chol, 7-OH-Chol, and 7-keto-Chol decreased or did not change significantly in *Dhcr7*^-/-^ cortices relative to *Dhcr7*^+/+^ cortices.

**Figure 2.**
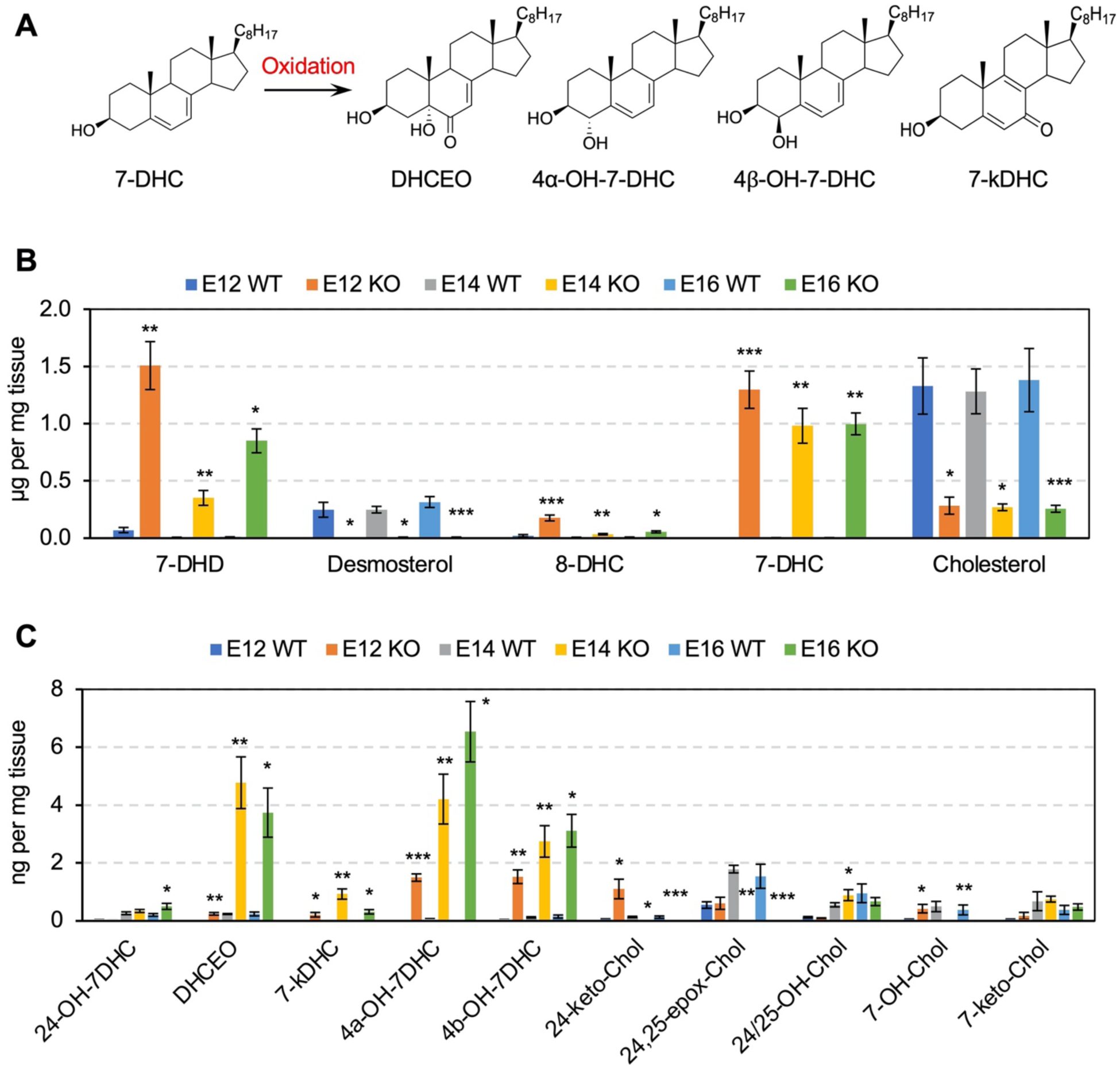
Cholesterol precursor 7-DHC and 7-DHC-derived Oxysterols are accumulated in *Dhcr7^-/-^* mouse embryonic cortex. (A) Chemical structures of 7-DHC-derived oxysterols. LC-MS/MS analysis of (B) cholesterol and its precursors and (C) 7-DHC-derived oxysterols in *Dhcr7*^+/+^ and *Dhcr7*^-/-^ embryonic cortex during development. Error bars indicate standard deviation. *, *p* < 0.05; **, *p* < 0.005; ***, *p* < 0.001; n = 3 biological replicates per group. **Figure 2-Figure Supplement 1.** Cholesterol precursor 7-DHC and 7-DHC-derived Oxysterols are accumulated *in* SLOS-derived human iPSCs and NPCs.

Furthermore, significant accumulation of 7-DHC and 7-DHC-derived oxysterols and reduction of cholesterol were also found in both SLOS patient-derived hiPSCs and NPCs relative to wild type (Figure 2-Figure Supplement 1). Taken together, *DHCR7* mutations led to accumulation of 7-DHC and 7-DHC-derived oxysterols in murine and human cortical precursors.

### Knockdown (KD) of Dhcr7 Causes Increased Neurogenesis and Depletion of Cycling Precursors in Murine and Human NPC cultures

To examine a potential role of *Dhcr7*, we generated three *Dhcr7* short hairpin RNAs (shRNAs)-EGFP reporters and transfected them into 293T cells along with murine *Dhcr7* cDNA-expressing plasmids. We found that *Dhcr7* shRNA2 was the most effective among those shRNAs and was chosen to examine *Dhcr7* function during neurogenesis from E12.5 cortical precursors in culture (Figure 3A). When cortical precursors were transfected with *Dhcr7* shRNA2-EGFP, the shRNA significantly decreased the percentage of transfected cells expressing *Dhcr7* (Figure 3B-C). *Dhcr7* KD led to a significant decrease in proliferation of cortical precursors as measured by Ki67 immunostaining and significant increase of βIII-tubulin+ neurons 3 days post-transfection, but it did not affect cell survival as examined by immunostaining for CC3 (Figure 3D-G). The significant decrease of precursor proliferation was caused by a decrease in Pax6+ radial precursors (Figure 3H).

**Figure 3.**
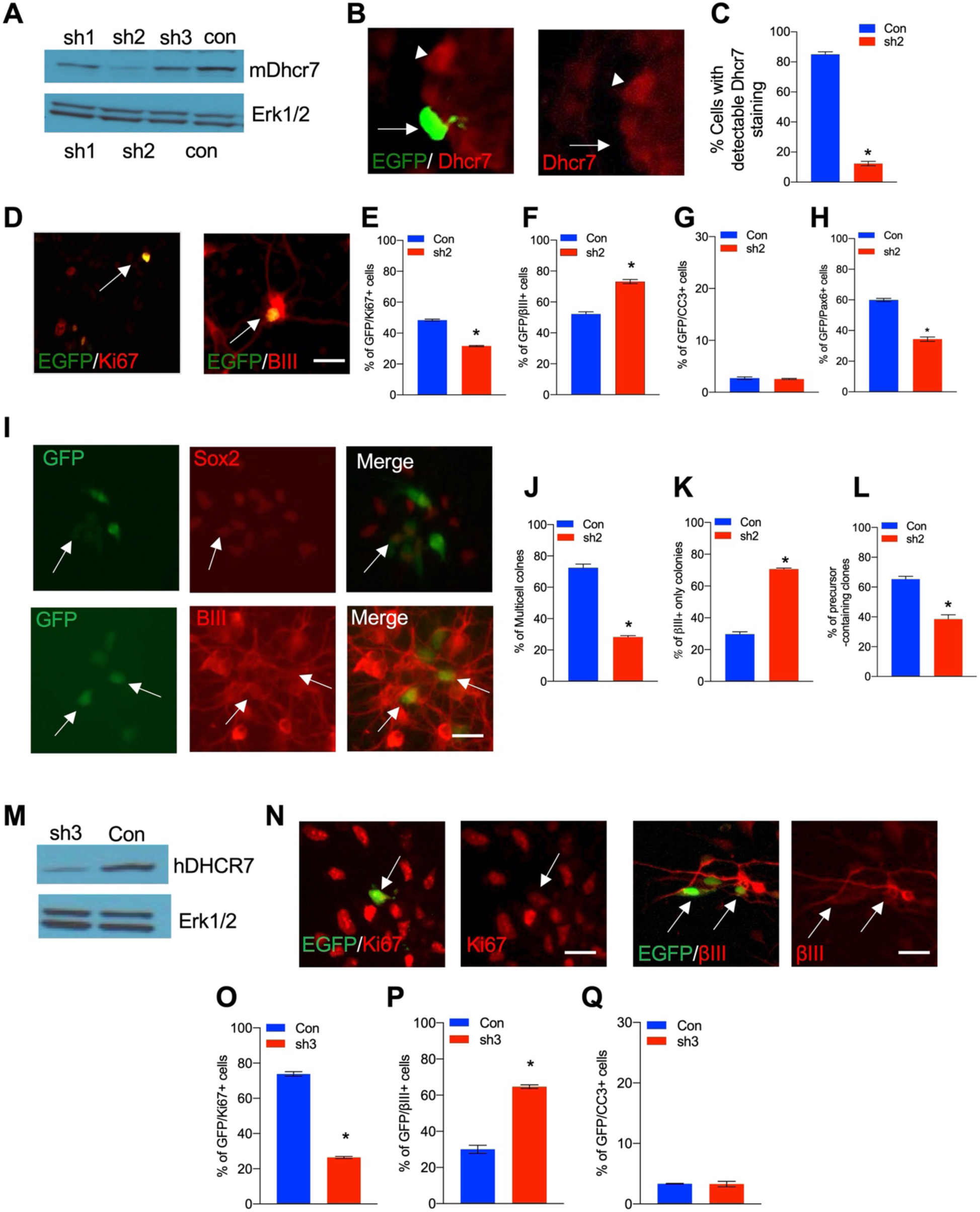
Knockdown of Dhcr7 causes increased neurogenesis and depletion of cycling precursors in murine and human NPC cultures. (A) Western blot for Dhcr7 in 293T cells transfected with control or individual murine *Dhcr7* shRNAs. The blot was re-probed for Erk1/2 as a loading control. (B and C) Mouse cortical precursors were transfected EGFP-*Dhcr7* shRNAs (sh2) or EGFP-control (Con) and immunostained for EGFP and Dhcr7 (red) 2 days later and EGFP+ cells expressing detectable Dhcr7 were quantified by fluorescence intensity (C). Arrow and arrowhead in (B) denote EGFP+/Dhcr7- and EGFP-/Dhcr7+ cells, respectively. (D-G) E12.5 cortical precursors were transfected with control or *Dhcr7* shRNAs and analyzed 3 days later. (D) Cultures were immunostained for EGFP (green) and Ki67 or βIII-tubulin (red; double-labelled cells in orange are indicated with arrows) or CC3 and the proportion of total EGFP+ cells that were also positive for Ki67 (E), βIII-tubulin (F) or CC3 (G) was quantified. *, *p* < 0.001; n = 3. Scale bar = 50 μm. In all cases, error bars denote SEM. (H-K) E12.5 precursors were co-transfected with the PB transposase and PB-EGFP-control (Con) or PB-EGFP-*DHCR7* shRNA (sh2). (H) Cultured cells were immunostained for EGFP (green), Sox2 (red) and βIII-tubulin (red) after 3 days and quantified for clones greater than one cell in size (I), neuron-only clones (J), and clones with at least one Sox2+ precursors (K). Arrows in (I) top denote EFGP+/ Sox2+ precursors. Arrow in (I) bottom denote EGFP+/βIII-tubulin+ cells. *, *p* < 0.001; n = 3. (M) Western blots of DHCR7 in 293T cells transfected with human control (Con) or human-specific *DHCR7* shRNA (sh3) plus human *DHCR7*-expressing plasmid, analyzed after 24 hr. The blot was re-probed for Erk1/2. (N) Human cortical precursors were transfected with EGFP-control (Con) or EGFP-*DHCR7* shRNA (sh3). Cells were immunostained 3 days later for EGFP (green) and Ki67 (red), βIII-tubulin (red) or CC3 and the proportion of total EGFP+ cells that were also positive for Ki67 (O), βIII-tubulin (P), or CC3 (Q) was quantified. *, *p* < 0.001; n = 3. Scale Bar = 50 μm. Allows in (N) denote double-positive cells. **Figure 3-Figure Supplement 1.** Rescue of the neurogenesis phenotype in *Dhcr7*-knockdown mouse cortical precursors by human *DHCR7* cDNA expression vector.

*Dhcr7* KD also decreased self-renewal of radial precursor as demonstrated by clonal analysis with piggybac (PB) transposon, which permanently labels precursors and their progeny. Cortical precursors were transfected with PB transposase and PB-*Dhcr7* shRNA2-EGFP or control shRNA-EGFP, and cultures were immunostained 3 days post-transfection for EGFP, the precursor markers, Sox2, and βIII-tubulin (Figure 3I-L). KD of *Dhcr7* reduced EGFP+ multicellular clones (Figure 3J) whereas neuron-only (βIII-tubulin+) clones were increased (Figure 3K). Furthermore, the number of precursors in clones containing at least one Sox2+ cells are decreased (Figure 3L).

To ensure that these changes were *Dhcr7* shRNA-dependent, we performed a rescue experiments using human *DHCR7* cDNA that is resistant to the murine *Dhcr7* shRNA (Figure 3-Figure Supplement 1). The murine shRNA did not affect the expression of human DHCR7 cDNA as confirmed by co-transfection of the murine shRNA and human *DHCR7* cDNA in 293T cells and western blot analysis 2 days later (Figure 3-Figure Supplement 1; Panel A). Furthermore, precursors were co-transfected with murine *Dhcr7* shRNA-EGFP or control shRNA-EGFP +/− human *DHCR7* cDNA and immunostained 3 days later for Ki67 or βIII-tubulin (Figure 3-Figure Supplement 1). The human *DHCR7* cDNA showed significant rescue of the murine *Dhcr7* shRNA KD phenotypes, confirming the specificity of the shRNA.

Finally, we asked whether *Dhcr7* also plays a role in human iPSC-derived NPCs (Figure 3M-Q). To examine the function of DHCR7, we generated a human-targeted *DHCR7* shRNA-EGFP and confirmed its efficiency by transfecting it into 293T cells along with human DHCR7 expressing vectors (Figure 3M). Human NPCs were transfected with this shRNA-EGFP and analyzed by immunostaining 4 days later, which showed that the KD of human *DHCR7* reduced EGFP+ and Ki67+ proliferating precursors whereas increased EGFP+ and βIII-tubulin+ newborn neurons (Figure 3N-Q), similar to those observed in murine cortical precursors and SLOS hNPCs.

### 7-DHC Derived Oxysterols Lead to Similar Neurogenic Defects as loss of Dhcr7 in Murine Cortical precursors in vitro

Oxysterols have been found to influence biological processes, including proliferation, differentiation and cell survival (DuSell et al., 2008; Theofilopoulos et al., 2013; Voisin et al., 2017). Thus, we asked whether 7-DHC-derived oxysterols play regulatory roles in cortical precursor biology. To assess potential effect of these 7-DHC-derived oxysterols on cortical precursor behaviors, cortical precursors from E12.5 wild-type embryos were cultured and treated with different concentrations of individual oxysterols for 3 days, followed by immunostaining for Ki67, βIII-tubulin, and CC3 (Figure 4A). Notably, DHCEO-treated cortical precursor showed significant increase in the proportion of βIII-tubulin+ newborn neurons and significant decrease in the proportion of Ki67+ precursors (Figure 4B-C). This effect of DHCEO was observed in a dose-dependent manner up to 3.5 µM, which is roughly the concentration of DHCEO in the brains of P0 *Dhcr7*-KO mice (Xu et al., 2011). There was no significant change in cell death/survival with DHCEO treatment (Figure 4D). The other oxysterols showed notable toxic effect at a concentration of 5 µM as indicated by increased CC3+ cells (Figure 4G, 4J, and 4M). Note that the level of 7k-DHC is the lowest among the oxysterols, only reaching 0.9 ng/mg of tissue (equivalent to 2.3 µM), so we do not expect the highest concentration examined, 5 µM, for this oxysterol is relevant to the neurogenic phenotype. Interestingly, 4*α*-OH-7DHC treatment at a concentration of 2 µM also displayed significant increases in βIII-tubulin+ neurons and decreases in Ki67+ precursors without significant changes in cell death/survival (Figure 4H-J). However, at 5 µM, CC3+ cells were significantly elevated at the expense of βIII-tubulin+ neurons (Figure 4J). To summarize, treatment with DHCEO at physiologically relevant concentrations completely replicated the premature neurogenic phenotype observed in murine and human SLOS NPCs while 4*α*-OH-7DHC may also contribute to the phenotype despite its toxicity at high concentration.

**Figure 4.**
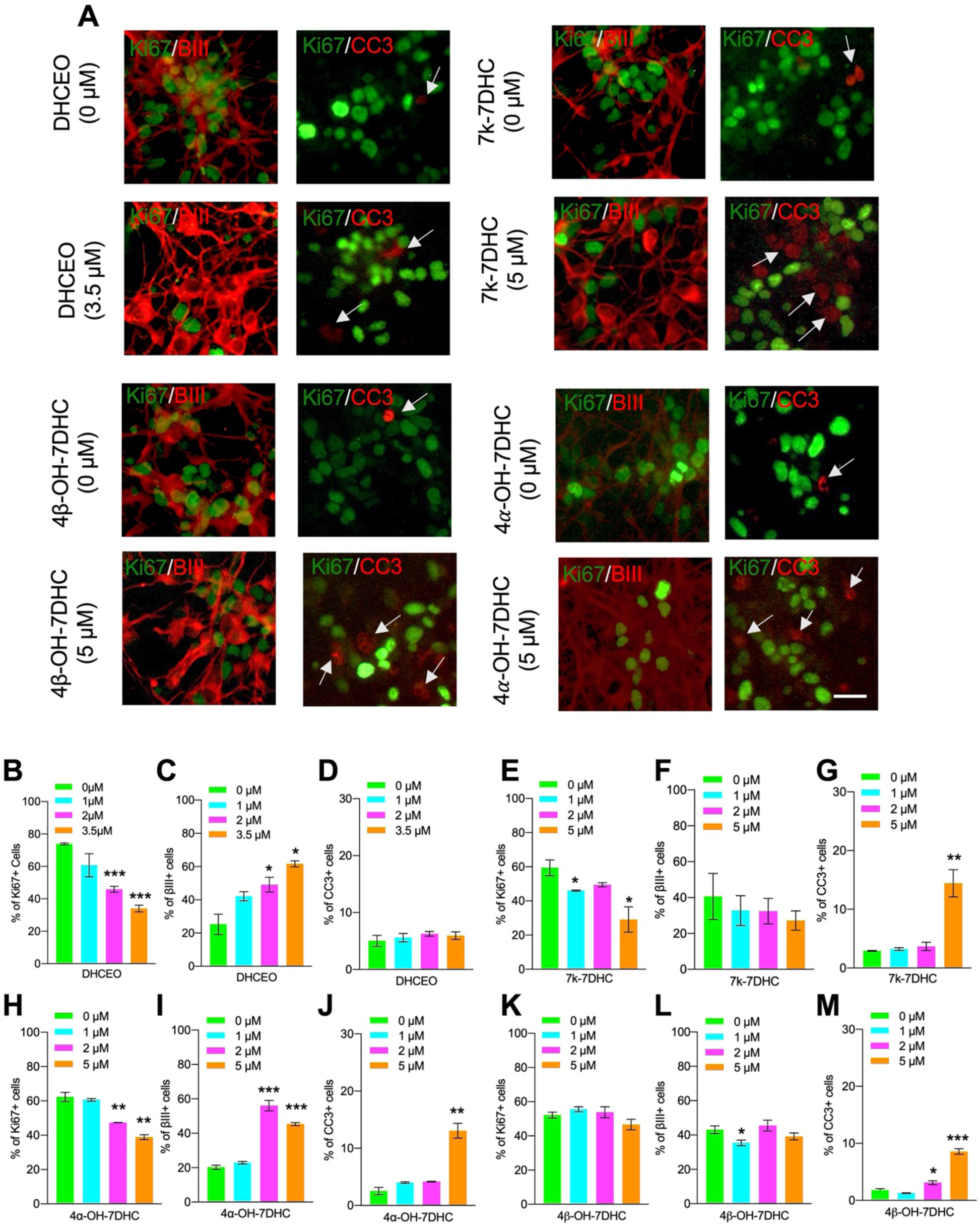
7-DHC Derived Oxysterols Lead to Similar Neurogenic Defects as loss of Dhcr7 in Murine Cortical precursors in vitro. (A-M) E12.5 cortical precursors were cultured for 2 days in the presence of different concentrations of 7-DHC-derived oxysterols and quantified. Cell were immunostained for Ki67 (green), βIII-tubulin (red), and CC3 (red, arrow) after 3 days and the proportions of Ki67+ (B,E,H,K), βIII-tubulin+ (C,F,I,L), and CC3+ (D,G,J,M) cells were determined. Error bars indicate SEM. *, *p* < 0.05; **, *p* < 0.005; ***, *p* < 0.001. n = 3 per experiment. Scale Bar = 50 μm.

### Loss of Dhcr7 causes premature differentiation and alter neocortical cytoarchitecture in vivo

We asked whether loss of *Dhcr7* disrupted cortical development *in vivo* as was observed *in vitro*. To examine this, cortical sections from *Dhcr7*^-/-^ mice at E18.5 were immunostained for the expression of well-defined cortical layer markers: the layer 2-4 marker Brn2, the layer 5 marker CTIP2, and the layer 6 marker Tbr1 (Molyneaux et al., 2007) (Figure 5A). The analyses showed loss of *Dhcr7* led to reduction of all cortical layers, resulting in reduction of overall size of the neocortex (Figure 5B-C). To further evaluate the effects of *Dhcr7* mutations in fate decision of cortical precursors during corticogenesis, we immunostained cortical sections from *Dhcr7*^-/-^ mice at E14.5 and E15.5 for Tbr1 and Satb2, which label early-born and late-born cortical neurons, respectively (Figure 5D-E). The analyses showed the significant increase of Tbr1+ neurons and Satb2+ neurons in cortical section from *Dhcr7*^-/-^ mice at E14.5 and E15.5, respectively (Figure 5F-G). This aberrant increased production of neurons with loss of *Dhcr7* potentially hinders proper localization of developing neurons in the cortex. To test this hypothesis, we injected pregnant dams with 5-ethynyl-2-deoxyuridine (Edu) at E12.5 to label proliferating radial precursors and analyzed *Dhcr7*^+/+^ and *Dhcr7*^-/-^ littermates three days later at E15.5 to identify the locations of Edu-labeled cells in the cortex (Figure 5H-I). Majority of Edu-labeled cells migrated into the cortical plate in the KO mice whereas Edu-labeled cells were scattered throughout the intermediate zone (IZ) and the cortical plates in the wild-type littermates. These observations suggest premature increases in precursor differentiation, which raises the possibility of activating major neurogenic signaling pathways during corticogenesis. RTKs and their downstream targets are known to play important roles in precursor proliferation and differentiation in developing cortex, and TrkB, one of the major RTKs expressed in cortical precursors, regulates proliferation and differentiation into neurons by activating the MEK-ERK-C/EBP pathway (Barnabe-Heider and Miller, 2003; Bartkowska et al., 2007; Menard et al., 2002). Thus, we examined the activity of TrkB-MEK-ERK-C/EBP pathway by western blot with antibodies for phosphorylated activated MEK1 and C/EBPβ at E15.5 of *Dhcr7*^-/-^ cortices (Figure 5J). This analysis showed that relative to total MEK1 and ERK1/2 levels, their phosphorylated forms increased in *Dhcr7*^-/-^ cortices. This observation suggests that loss of *Dhcr7* can dysregulate an RTK-dependent signaling pathway that is critical for neural precursor proliferation and differentiation.

**Figure 5.**
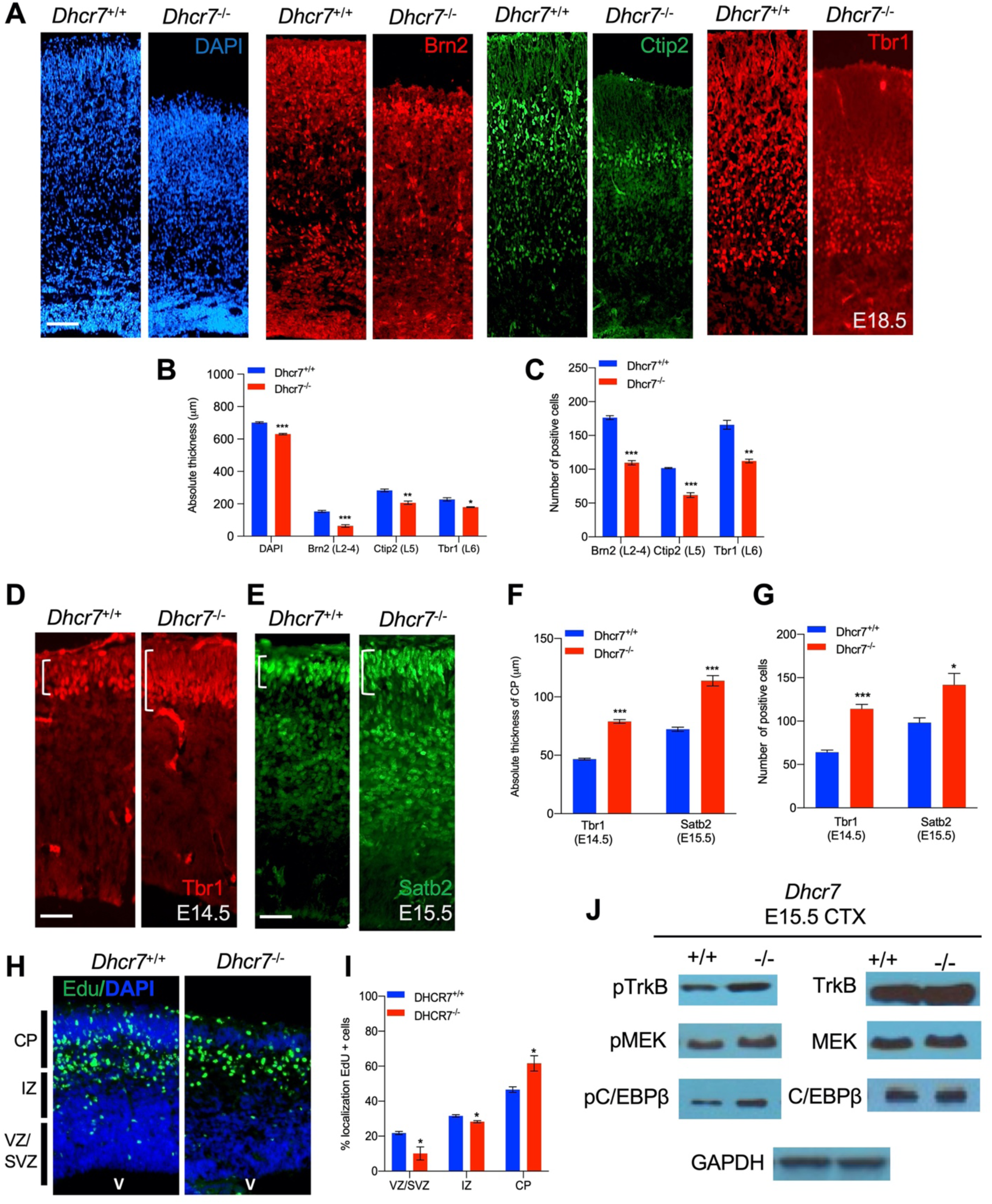
*Dhcr7^-/-^* mice display premature neurogenesis, altered neocortical cytoarchitecture *in vivo*, and increased activity of TrkB neurogenic signaling pathway. (A) E18.5 cortical sections from *Dhcr7*^+/+^ and *Dhcr7*^-/-^ were immunostained for Tbr1 (red), Ctip2 (green) and counterstained with DAPI (blue). (B and C) Quantifications of absolute thickness and number of positive cells (C). (D-G) Cortical sections from E15.5 *Dhcr7*^+/+^ and *Dhcr7*^-/-^ mice were immunostained for Satb2 (D, green) and Tbr1 (E, red). Quantifications of absolute thickness (F) and number of positive cells (G) for Satb2 and Tbr1. (H) E15.5 cortical sections from *Dhcr7*^+/+^ and *Dhcr7*^-/-^ embryos EdU-labeled at E12.5 were immunostained for EdU (green) and counterstained with DAPI (blue). (I) Quantification of relative location of EdU+ cells in cortical sections. (J) E15.5 cortices were isolated from *Dhcr7*^+/+^ and *Dhcr7*^-/-^ embryos and analyzed by western blot for phospho-TrkB, phospho-MEK, or phospho-C/EBPβ. Blots were re-probed with antibodies for total GR, TrkB, MEK, C/EBPβ and GAPDH as loading controls. Error bars indicate SEM. *, *p* < 0.05; **, *p* < 0.005; ***, *p* < 0.001. n = 3 per experiment. Scale Bar = 50 μm.

The above observations indicate premature precursor differentiation at the expense of depleting the proliferative precursor pool. To examine this hypothesis, pregnant dams were injected with Edu at different time points of cortical development, and *Dhcr7*^+/+^ and *Dhcr7*^-/-^ littermates were analyzed 18 hours later to quantify the number of cycling progenitors that have exited the cell cycles (Figure 6A-B). Cells that had left the cell cycle were stained as Edu+ and Ki67-whereas cells that re-entered the cell cycle were stained as Edu+ and Ki67+. The index of the cell cycle exit is defined by the percentage of Edu+/Ki67-cells in total Edu+ cells. The index showed that the number of cells exiting the cell cycle in *Dhcr7*^-/-^ cortices was significantly increased at E13.5 and E14.5 compared with *Dhcr7*^+/+^ cortices, indicating that loss of *Dhcr7* increased cell cycle exit. Increased cell cycle exit of cortical precursors can potentially arise from changes in proliferation. To test this possibility, pregnant dams were pulsed with Edu for 2 hours to label all cells in S-phase and the fraction of cortical precursors in S-phase, a proliferating index from E13.5 to E15.5 in *Dhcr7*^-/-^ and *Dhcr7*^+/+^ cortices, was quantified (Figure 6C-D). The proliferating index was determined by the percentage of Edu+ and Ki67+ cells out of total Ki67+ cells, which provides an estimate of cell cycle length because the length of S phase remains relatively constant in mammalian cells, whereas the length of G1 phase regulates proliferation (DiSalvo et al., 1995). The analyses showed a significant decrease in the proliferating index in *Dhcr7*^-/-^ cortices compared to *Dhcr7*^+/+^ cortices from E13.5 to E15.5 (Figure 6C-D), indicating slower cycle progression and longer cycle length in *Dhcr7*^-/-^ cortices (Chenn and Walsh, 2002). Indeed, this accelerated depletion of the progenitor pool in *Dhcr7*^-/-^ cortices result in decreases in the size of VZ/SVZ compared to *Dhcr7*^+/+^ cortices from E13.5 to E15.5 (Figure 6E-F). Taken together, these results suggest that loss of *Dhcr7* leads to a decrease in the overall number of cortical precursors with increased cell cycle exit and decreased proliferation index via activation of RTK-mediated MEK-ERK-C/EBP pathway.

**Figure 6.**
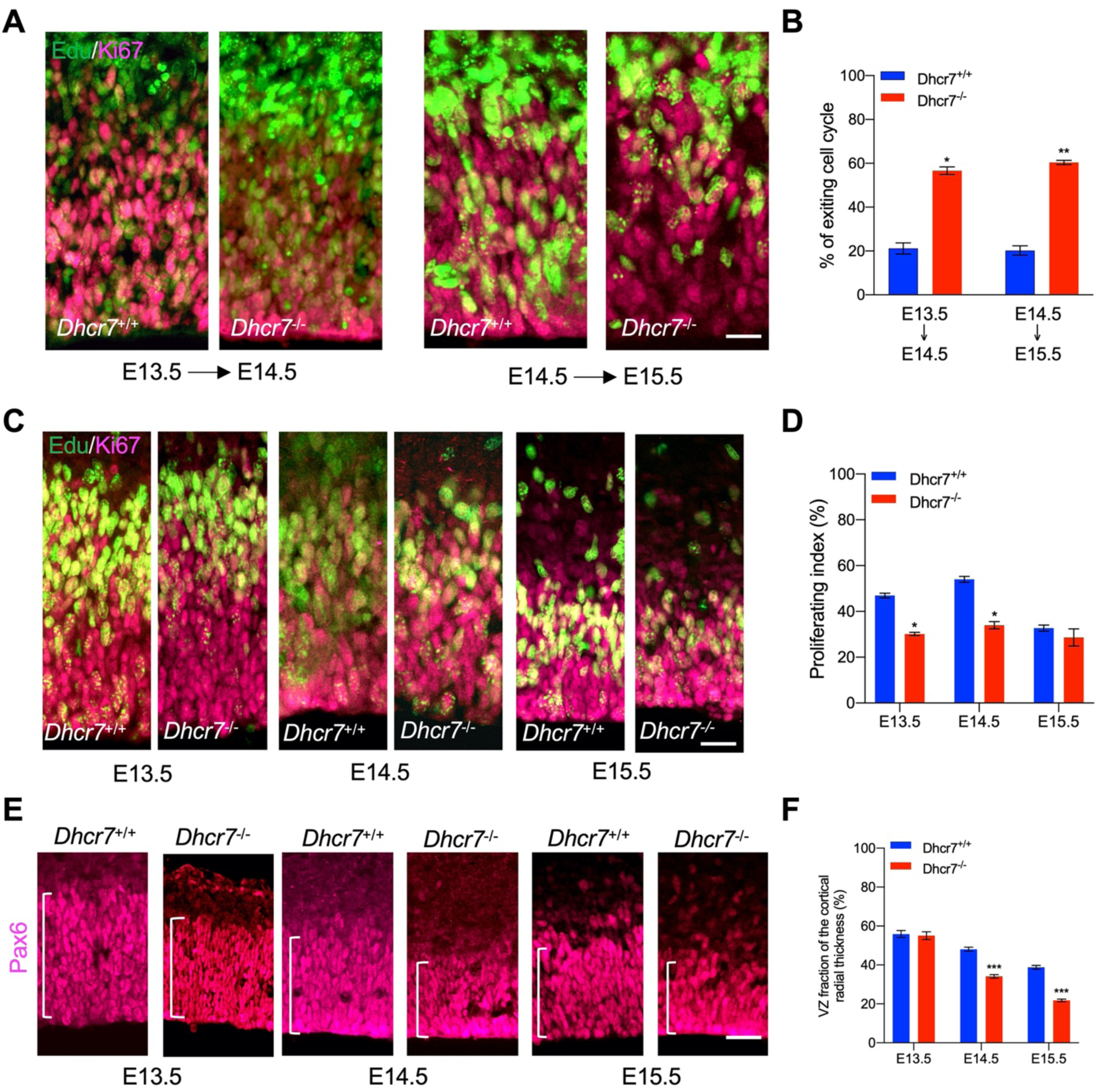
*Dhcr7-/-* mice show accelerated cell cycle exiting and depletion of cortical precursor cells *in vivo*. (A) Cortical sections from *Dhcr7*^-/-^ and *Dhcr7*^+/+^ embryos labeled by Edu at different developmental stages were immunostained 18 hours later for Edu (green) and Ki67 (Magenta). (B) Quantification of cell-cycle exit index of *Dhcr7*^-/-^ cortices compared with *Dhcr7*^+/+^ cortices. (C) Cortical sections from *Dhcr7*^-/-^ and *Dhcr7*^+/+^ embryos labeled by Edu injection at different developmental stages were immunostained 2 hours later for Edu (green) and Ki67 (Magenta). (D) Quantification of proliferation index of *Dhcr7*^-/-^ cortices compared with *Dhcr7*^+/+^ cortices. (E) Coronal cortical sections immunostained for Pax6 cortical precursor marker at different developmental stages. (F) Quantification of the relative size of the Pax6+ region shown as fractions of the whole cortical radial thickness. Error bars indicate SEM. *, *p* < 0.05; **, *p* < 0.005; ***, *p* < 0.001. n = 3 per experiment. Scale Bar = 50 μm.

### Perturbation in DHCR7 Causes Gene Expression Changes in Neurogenic Pathways in Human NPCs

To ask whether *DHCR7* mutations influence the NPC biology, we aim to compare gene expression profiles between hNPCs with *DHCR7* mutations versus hNPCs from healthy individuals. To do so, we expanded the hNPCs derived from hiPSCs in the presence of essential growth factors and carried out RNA sequencing (RNAseq) on these cells. Relative to wild-type hNPCs, 2357 genes were significantly differentially expressed in SLOS hNPCs (1072 upregulated and 1285 downregulated genes with adjusted *p* value < 0.05), distinguishing the SLOS and wild-type precursors (Figure S1A; Table S6 for RNAseq data). Gene ontology and pathway analysis found that PI3K-Akt signaling pathway, MAPK signaling, and Ras signaling pathways are among those with the highest *p*-values (Figure S1B-C; Table S5). Notably, MAP2K1(also known as MEK1) was found to be upregulated in SLOS patient-derived hNPCs. MAP2K1 is a key signaling molecule in MAPK and Ras signaling pathways, which are known to play important roles in precursor proliferation and differentiation (Bonni et al., 1999; Menard et al., 2002; Yang et al., 2013), consistent with the phenotypes that were observed in *Dhcr7*-KO mice and SLOS patient-derived hNPCs *in vitro*.

### 7-DHC-derived oxysterol, DHCEO, activates cortical neurogenesis via activation of glucocorticoid receptor and inhibition of GR or inhibition of the formation of DHCEO rescues the neurogenic defects in SLOS NPCs

As shown in Figure 4, treatment of WT mouse NPCs with 7-DHC-derived oxysterols, particularly DHCEO, can replicate the same aberrant premature neurogenesis observed in SLOS NPCs. However, the mechanism of action of DHCEO remains unknown. Interestingly, 6-oxo-cholestan-3β,5*α*-diol (OCDO), a cholesterol-derived oxysterol that is structurally similar to DHCEO, is shown to bind and activate glucocorticoid receptor (GR) (Voisin et al., 2017) (Figure 7-Figure Supplement 1; Panel A). Additionally, GR activation is shown to activate TrkB (Jeanneteau et al., 2008), which leads to further activation of RTK-mediated MEK-ERK pathway (Barnabe-Heider and Miller, 2003; Bartkowska et al., 2007; Menard et al., 2002). TrkB and RTK-mediated MEK-ERK pathway are necessary for neurogenesis during embryonic cortical development. Thus, we asked whether DHCEO can activate the GR and further lead to the activation of MEK-ERK neurogenic pathway. To investigate this, we first examined if GR was phosphorylated and activated in the embryonic cortices from *Dhcr7*^-/-^ and *Dhcr7*^+/+^ embryos at E15.5 in addition to TrkB, MEK, and C/EBPβ shown in Figure 5J. Western blots demonstrated that phosphorylation of GR was indeed increased in *Dhcr7*^-/-^ cortices comparing to *Dhcr7*^+/+^ cortices (Figure 7A).

**Figure 7.**
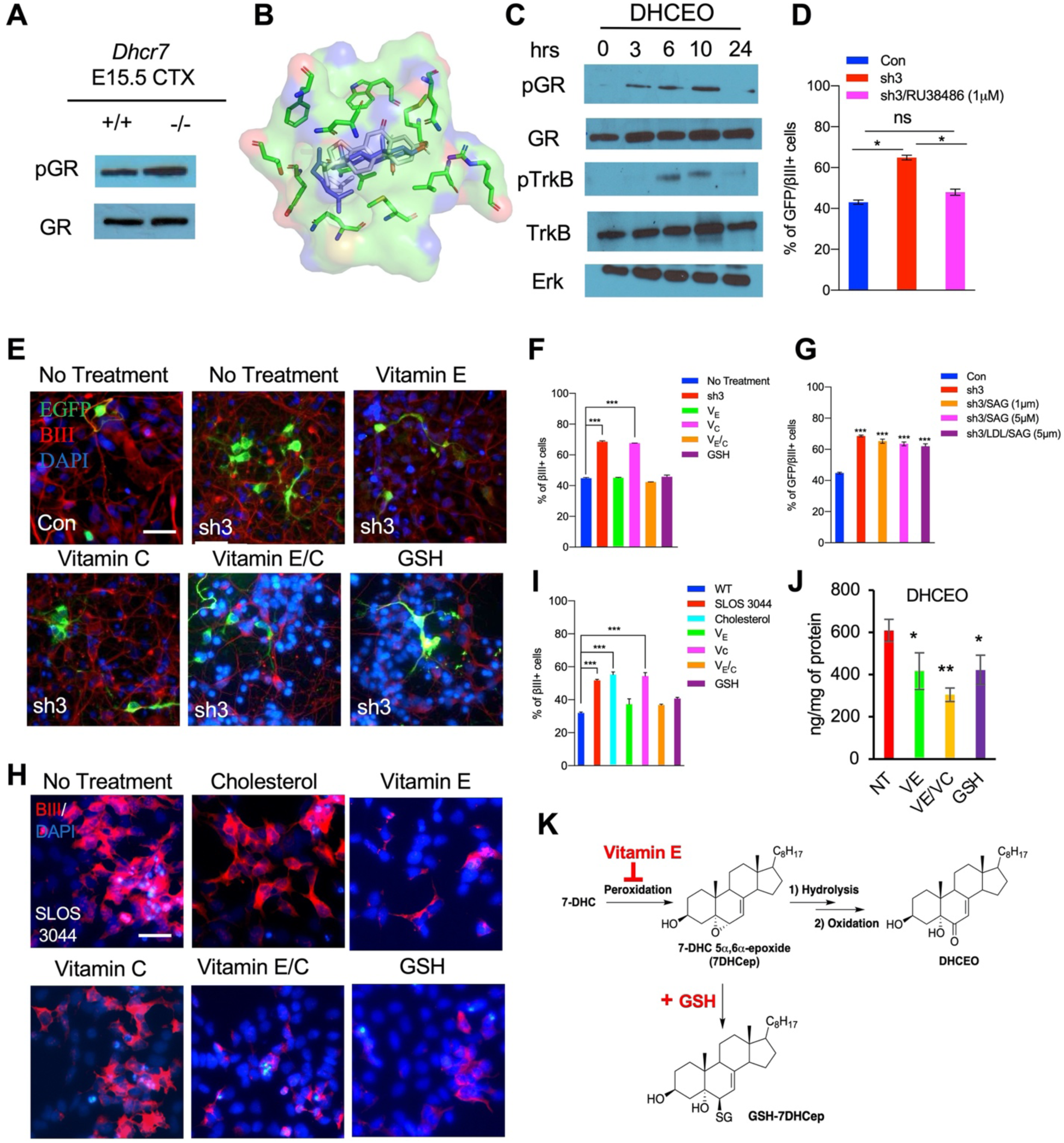
DHCEO activates cortical neurogenesis via activation of glucocorticoid receptor and inhibition of the effect or the formation of DHCEO rescues the neurogenic defects in SLOS NPCs. (A) Western blot showing increased phospho-GR in E15.5 *Dhcr7*^-/-^ mouse brain relative to *Dhcr7*^+/+^. (B) Image of the docked position of DHCEO (white) and OCDO (blue) in the ligand binding pocket of GR. (C) Human neural progenitors were treated with 3.5 μM DHCEO over the indicated time periods. Lysates were probed with phosphor-GR and phosphor-TrkB and re-probed with antibodies for total GR, total TrkB or total ERK as loading controls. (D) Human NPCs were transfected with EGFP-control (Con) or EGFP-*DHCR7* (sh3) shRNA. Cells were treated with 1 μM RU38486, a selective GR antagonist one day after transfection. Three days post-transfection, cells were immunostained for EGFP and βIII-tubulin and quantified. (E) Wild type hNPCs were transfected with EGFP-control (Con) or EGFP-*DHCR7* shRNA (sh3), and then treated with vitamin E, vitamin C, vitamin E/C, or glutathione (GSH). Three days post-transfection, cells were immunostained for EGFP and βIII-tubulin. (F,G) Quantification of EGFP and βIII-tubulin+ cells in *DHCR7*-KD wild-type hNPCs treated with various antioxidants, SAG, or LDL+SAG. (H) SLOS hNPCs were treated with cholesterol or various antioxidants and were immunostained for βIII-tubulin and DAPI. (I) Quantification of the proportion of βIII-tubulin+ cells in wild type hNPCs, and SLOS hNPCs treated with cholesterol or various antioxidants. (J) Quantification of DHCEO by LC-MS/MS in SLOS hiPSCs treated with vitamin E, vitamin E/C, and GSH. (K) Proposed mechanisms of actions of vitamin E and GSH during free radical oxidation of 7-DHC. Error bars indicate SEM. *, *p* < 0.05; **, *p* < 0.005; ***, *p* < 0.001. n = 3 per experiment. Scale Bar = 50 μm. **Figure 7-Figure Supplement 1.** Antioxidants rescue the neurogenic phenotype in human and murine NPCs with *Dhcr7* mutations.

To ask if DHCEO could physically interact with GR, we performed molecular docking simulation between DHCEO and the ligand binding domain of GR. Molecular docking is an effective computational approach to understand protein-ligand interaction between small molecules and receptor proteins both energetically and geometrically (Ferreira et al., 2015). As seen in Figure 7B, DHCEO docks in the binding pocket of human GR favorably with a docking score of −9 (a negative value suggests favorable interactions), comparable to the docking score of OCDO at −9.5.

We further examined whether DHCEO could initiate GR and TrkB activation in wild-type hNPCs *in vitro*. Cultured hNPCs were exposed to DHCEO (3.5 μM), a physiologically relevant concentration, harvested at the different time points (3, 6, 9 and 24 hrs), and evaluated for GR and TrkB activation by western blots (Figure 7C), which revealed that DHCEO-treated hNPCs showed a gradual increase of phosphorylated GR starting at 3 hrs and peaking at 10 hrs of exposure. As GR phosphorylation increased, DHCEO-treated hNPCs also showed the phosphorylation of TrkB from 6 hrs to 10 hrs exposure, which dissipated at 24hrs exposure. As no exogenous GR ligands and neurotrophins were added in these experiments, these results indicated that DHCEO caused the increased phosphorylation of GR, which led to the activation of TrkB phosphorylation in human NPCs *in vitro*.

To confirm that GR activation could cause aberrant premature neurogenesis, we cultured human cortical precursors and transfected them with *hDHCR7* shRNA-EGFP and control shRNA-EGFP. 12 hours post-transfection, a glucocorticoid receptor antagonist, RU38486, or vehicle control was added and cultured for 3 days. Cultures were immunostained for EGFP and βIII-tubulin to evaluate the effects on neurogenesis. The analysis showed that inhibition of GR by RU38486 rescued the *DHCR7*-KD-mediated increase in neurogenesis down to the level observed in the control shRNA (Figure 7D).

Finally, we assessed whether aberrant premature neurogenesis was due to DHCEO, as it is a major oxysterol derived from free radical (non-enzymatic) oxidation of 7-DHC (Xu et al., 2011) (Figure 2). Antioxidants can prevent or slow down formation of oxysterols via the free radical mechanism and protect cells from deleterious effects of oxysterols (Yin et al., 2011). In particular, vitamin E (V_E_), but not vitamin C (V_C_), has been shown to effectively inhibit formation of 7-DHC derived oxysterols in SLOS patient-derived fibroblasts (Korade et al., 2014). Combination with V_C_, which recycle oxidized V_E_, enhances the antioxidant activity of V_E_ against lipid peroxidation (Doba et al., 1985). Furthermore, glutathione (GSH) is an important detoxifying molecule that is abundant (in mM) in cells and can potentially inhibit the formation of DHCEO by reacting with its electrophilic precursor, 7-DHC 5*α*,6*α*-epoxide (Porter et al., 2020; Xu et al., 2011). To test this possibility, we treated *hDHCR7* KD human cortical precursors with antioxidants (V_E_, V_C_, V_E_/V_C_, or GSH) and then immunostained for EGFP and βIII-tubulin to evaluate the effects on neurogenesis (Figure 7E-F). The analysis indicated that V_E_, V_E_/V_C_, or GSH, but not V_C_ alone, effectively reversed increased neurogenesis caused by *hDHCR7* KD to the level observed in the control. Potential detoxifying effects of these antioxidants were further tested in human SLOS patient-derived NPCs as well as cortical precursors from *Dhcr7*^-/-^ mouse embryos *in vitro* (Figure 7H-I and Figure 7-Figure Supplement 1, panels B-E). Treatments with V_E_, V_E_/ V_C_ and GSH were again found to effectively reduce the increased neurogenesis in both human and murine cortical precursors with *Dhcr7* mutations down to the level observed in WT samples. LC-MS/MS analysis confirmed that V_E_, V_E_/V_C_, or GSH indeed inhibited the levels of DHCEO in SLOS hiPSCs (Figure 7J). These results suggest that free radical chain-breaking antioxidant, such as V_E_, and nucleophilic antioxidant, such as GSH, can effectively rescue the neurogenesis phenotype observed in SLOS NPCs by inhibiting the free radical oxidation of 7-DHC at different stages (Figure 7K).

In addition to antioxidant treatment, we tested whether restoration of cholesterol could potentially rescue aberrant premature neurogenesis observed in both human and murine SLOS NPCs (Figure 7I, S6C and S6E). Low-density lipoprotein was added to cultures to examine the effects of cholesterol on neurogenesis. The results revealed that addition of cholesterol did not prevent increased neurogenesis in SLOS NPCs. Because DHCEO has been shown to inhibit Smo in the Hh signaling pathway (Sever et al., 2016), we also tested whether supplementation of SAG (a Smo agonist) can rescue the phenotype with or without cholesterol supplementation in *DHCR7*-KD hNPCs (Figure 7G). However, none rescued the neurogenesis phenotype. Thus, the lack of cholesterol and inhibition of Hh signaling do not contribute to the neurogenesis phenotype observed in SLOS NPCs.

Taken together, the abnormal premature neurogenesis in cortical precursor with *Dhcr7* mutations was caused by DHCEO activation of GR and further activation of Trk-mediated neurogenic signaling pathway, but not by the deficiency in cholesterol, and inhibition of the formation of DHCEO or GR rescues the neurogenic defects.

## Discussion

Whether deficiency in cholesterol, accumulation of 7-DHC, or accumulation of 7-DHC oxysterols is more important for the neurological and developmental phenotypes observed in patients in SLOS remains unknown. Here, we characterized the neurogenic phenotype of SLOS cortical precursors *in vitro* and *in vivo* and showed that the accumulated 7-DHC-derived oxysterols, particularly DHCEO, disrupts normal embryonic neurogenesis by accelerating progressive depletion of cortical precursors through neurogenesis, ultimately resulting in greatly reduced cerebral cortex thickness with abnormal cortical layering.

The data presented here support four major conclusions. First, LC-MS/MS analyses indicate that *DHCR7* mutations promote significant accumulation of 7-DHC-derived oxysterols in murine and human embryonic cortical NPCs during cortical development. Second, our studies with KD or KO of *DHCR7* indicate that *DHCR7* is necessary for normal proliferation of embryonic NPCs and genesis of newborn neurons in culture and within the environment of the embryonic cortex. KD or KO of *DHCR7* leads to premature differentiation of cortical precursors to neurons and ultimately a perturbation in cortical cytoarchitecture. This occurs through the activation of neurogenic TrkB-MEK-C/EBP pathway. Third, we show that 7-DHC-derived oxysterols, especially DHCEO, display detrimental effects to normal neural development, and that DHCEO can activate GR, which is responsible for abnormal cortical precursor differentiation, since *DHCR7* KD/KO phenotype can be rescued by concurrent inhibition of glucocorticoid receptor. Fourth, we show that premature differentiation of cortical precursors with *DHCR7* mutations is cholesterol-independent since inhibition of 7-DHC-derived oxysterols via antioxidant treatment can rescue *DHCR7* KD/KO phenotype whereas the cholesterol supplementation cannot.

Based on these data, we propose a model where DHCEO binds and activates GR, and in doing so, controls the RTK-mediated neurogenic pathway, TrkB-MEK-C/EBP, and thus fate-decision of developing precursors and neurons. This model is consistent with previous studies showing that an oxysterol structurally similar to DHCEO, OCDO, can bind and regulate GR and its transcriptional activity (Voisin et al., 2017) and ligand-bound GR promoted TrkB phosphorylation, leading to activation of its downstream effectors, MEK and ERK (Jeanneteau et al., 2008). Our model of DHCEO exerting its effect on neurogenesis through GR does not contradict the previous finding by Francis et al., which suggests that Wnt/β-catenin inhibition contributes to precocious neurogenesis (Francis et al., 2016), because activation of GR has been shown to inhibit Wnt/β-catenin signaling in several systems (Olkku and Mahonen, 2009; Zhou et al., 2020).

This work represents the first comprehensive characterization of the neurogenic phenotype in SLOS mouse model and SLOS patient-derived NPCs. We demonstrate that the aberrant neural development observed in mice and human NPCs with *DHCR7* mutations occurs in a cholesterol-independent and 7-DHC-derived oxysterol-dependent manner. Successful rescue of the neurogenic phenotype by inhibiting GR or inhibiting the formation of oxysterols with antioxidants pave the wave for potential therapies toward the neurological defects observed in SLOS patients.

## Supporting information

Figure Supplements related to main figures

Differentially expressed genes; WT vs. SLOS hNPCs.

## Acknowledgements

This work was supported by a National Institutes of Health (NIH) grant R01HD092659 (LX) and a grant from Smith-Lemli-Opitz/RSH Foundation. JMH was supported by the University of Washington (UW) Environmental Pathology/ Toxicology Training Program (NIH T32 ES007032) and AL was supported by the UW Pharmacological Sciences Training Program (NIH T32 GM007750) and the Institute of Translational Health Sciences TL1 Program (NIH TL1 TR002318). We also thank Dr. Carol Ware and Christopher Cavanaugh for technical advices on hiPSC culture.

## Author Contributions

HT conceptualized, designed, performed, and analyzed most of the experiments and co-wrote the paper. LX conceptualized, designed experiments, analyzed data and co-wrote the paper. KMH and AL performed and analyzed LC-MS/MS. JMH analyzed RNA-sequencing experiments. DB performed and analyzed docking simulation.

## Competing interests

The authors declare no competing interests.

## Materials and Methods

### Animals

Animal use was approved by the University of Washington Institutional Animal Care and Use Committee. *Dhcr7* mice (B6.129P2(Cg)-*Dhcr7*^tm1Gst^/J) were maintained as heterozygous and were genotyped as described previously (Fitzky et al., 2001). The primers used are the following: *Dhcr7*-WT-F: 5’-GGATCTTCTGAGGGCAGCTT-3’; *Dhcr7*-WT-R: 5’-TCTGAACCCTTGGCTGATC-3’; Delta-Mut: 5’-CTAGACCGCGGCTAGAGAAT-3’. For *in vitro* transfection, C57BL/6J E12.5 pregnant mice were obtained by time-mating. Mating pairs of wild-type C57BL/6J mice were purchased from Jackson Laboratories (Bar Harbor, ME). All mice had free access to rodent chow and water in a 12-hour dark-light cycle room.

### Plasmids

The target sequence for murine *Dhcr7* shRNA was cloned into pLKO-UBI-GFP digested with EcoRI and PacI. pLKO-UBI-GFP was generated by digesting out hPGK promoter from pLKO.3G and ligating ubiquitin promoter (UBI) from pCLX-UBI-VenusN with PacI and BamHI. pLKO.3G was a gift from Christophe Benoist & Diane Mathis (Addgene plasmid # 14748; http://n2t.net/addgene:14748; RRID:Addgene_14748) and pCLX-UBI-VenusN was a gift from Patrick Salmon (Addgene plasmid # 27247; http://n2t.net/addgene:27247; RRID:Addgene_27247). cDNAs encoding murine (pCMV-mDHCR7-Myc/DDK) and human DHCR7 (pCMV-hDHCR7-Myc/DDK) were purchased from Origene (MR223420 and RC228922 for murine and human cDNA clone respectively). PB-EF1*α*-GreenPuro-H1MCS and Super PiggyBac transposase expression vector from System Biosciences (Cat#PBS506A-1 and Cat# PB210PA-1 respectively) were used for clonal analysis.

### Cortical precursor cell cultures

Murine cortical precursor cells were cultured as described previously (Barnabe-Heider and Miller, 2003; Tomita et al., 2020). Briefly, mouse cortical precursor cells from cortices were dissected from E12.5 wild-type or *Dhcr7*-KO mouse embryos in ice-cold HBSS (Invitrogen) and transferred into cortical precursor medium. The cortical precursor medium consisted of Neurobasal medium (Invitrogen) with 500 μM L-glutamine (Invitrogen), 2% B27 supplement (Invitrogen), 1% penicillin-streptomycin (Invitrogen) and 40 ng/ml FGF2 (BD Biosciences). The dissected tissue was mechanically triturated by a fire-polished glass pipette and plated onto 24-well plates coated with 2% laminin (BD Biosciences) and 1% poly-D-lysine (Sigma Aldrich). Plating density of the cortical precursors was 150,000 cells/well in 24-well plates for single embryo cultures and plasmid transfections.

### Human induced pluripotent stem cells (hiPSCs)

Human fibroblast cell lines (GM03044 and GM05788) isolated from patients with SLOS were obtained from Coriell Institute and reprogrammed as described elsewhere (Okita et al., 2011; Yu et al., 2007). Briefly, 1×10^6^ cells of human fibroblast cells at early passages were trypsinized (0.25% Trypsin/0.5mM EDTA, Gibco) and electroporated with indicated episomal plasmids using Amaxa Basic Nucleofector Kit for primary mammalian fibroblasts, program A-24 (Lonza). 1.0μg pSIN4-EF2-O2S and pSIN4-EF2-N2L were used for each electroporation. The electroporated human fibroblasts were seeded onto a gelatin-coated tissue culture dish and cultured with fibroblast medium (DMEM with 10% FBS, 2mM L-glutamine, 1% non-essential amino acids, 1% sodium pyruvate, and 1% penicillin and streptomycin) for 7 days and re-seeded onto feeder layer cells (irradiated SNL 76/7 feeder cells; a gift from Dr. Allan Bradley, Sanger Institute, UK) with hiPSC medium (DMEM/F12 medium with 20% knockout serum replacer, 1% sodium pyruvate, 1% non-essential amino acids, 0.007% 2-mercaptoethanol,1% penicillin and streptomycin, 0.1mM sodium butyrate, 50nM suberoylanilide hydroxamic acid, and 4ng/ml of bFGF) in 100mm tissue culture dishes and continued to be cultured until hiPSC colonies were visible (28 days from electroporation). The hiPSC colonies were then picked and further cultured for expansion.

### Maintenance and Differentiation of hiPSC

Undifferentiated hiPSCs were cultured and maintained in mTeSR Plus medium (StemCell Technologies) on Matrigel (BD Biosciences)-coated plates prior to the generation of human neural progenitor cells (hNPCs). hNPCs was generated using the STEMdiff SMADi Neural Induction kit (StemCell Technologies) according to the manufacturer’s protocol. Briefly, neuralized embryoid bodies (EBs) were generated by culturing small aggregates of hESCs in ultra-low attachment plates (Corning) in STEMdiff SMADi Neural induction medium. EBs were replated and cultured in poly-D-lysine/Laminin-coated plates with the neural induction medium until EBs formed neural rosette formations. Neural rosettes were collected and replated in poly-D-lysine/Laminin-coated plates for hNPCs outgrowth in neural induction medium until hNPCs were ready for the first passage. hNPCs were maintained in STEMdiff Neural Progenitor medium. For neurogenesis of hNPCs, hNPCs were cultured in Neurobasal A medium supplemented with B27 minus vitamin A, 1% penicillin-streptomycin and glutamax (all from Invitrogen) in poly-D-lysine/Laminin-coated plates (Konopka et al., 2012; Usui et al., 2017). For quantification, an average of 700 cells was counted per condition in 5-6 random fields per independent experiment.

### In vitro differentiation of iPSCs

For in-vitro differentiation of human iPSCs to the three germ layers, STEMdiff Trilineage Differentiation Kit (StemCell Technologies) was used as described by the manufacturer. Briefly, cells were harvested by Gentle Cell Dissociation Reagent (StemCell Technologies) and plated onto Matrigel-coated 24 well plates with mTeSR1 medium (StemCell Technologies). Cell density and viability were determined using trypan blue exclusion. Cells were seeded at a clonal density of 200,000 cell/cm^2^ for ectoderm and endoderm differentiation and 50,000 cell/cm^2^ for mesoderm differentiation. 24 hours after plating, cells were switched to the appropriate STEMdiff Trilineage medium for ectoderm, mesoderm and endoderm differentiation. Cells were cultured in the lineage specific medium for 5 days (mesoderm and endoderm lineages) and 7 days (ectoderm lineage) and were harvested and/or fixed for analyses of lineage-specific markers for the three germ layers (Figure 1-Figure Supplement 1).

### Transfection and quantification

For plasmid transfection of mouse cortical precursors, Lipofectamine LTX and Plus Reagent (Invitrogen) were used as described by the manufacturer. Briefly, 1 μg of DNA and 1 μl of Lipofectamine LTX and Plus Regent in 100 μl of Opti-MEM (both from Invitrogen) were mixed, incubated for 20 min, and added to precursors three hours after plating. For clonal analysis, 1.5 μg DNA, at 1:3 ratio of Super PiggyBac transposase expression vector to shRNA or control plasmids, were incubated as described above and added to cortical precursors three hours after plating. For plasmid transfection of human cortical precursors, Lipofectamine Stem Transfection Reagent (Invitrogen) was used as described by the manufacturer. Briefly, 1μg of DNA and 1μl of Lipofectamine Stem Transfection Reagent were prepared separately in 25μl of Opti-MEM medium (Invitrogen), mixed, incubated for 30 min and added to precursors one day after plating. The target sequences for murine and human *Dhcr7* shRNA were 5’-GGAAGGTGCTTCTTGTTTA-3’ and 5’-GGAAGTGGTTTGACTTCAA-3’ respectively. The target sequence for the control shRNA was 5’-TCCCAACTGTCACGTTCTC-3’. For quantification, immunostaining and image acquisition were performed, and > 100 cells per condition per experiment were counted and analyzed, and experiments were performed with 3 embryos per plasmid transfected and analyzed individually.

### Neurosphere cultures

E13.5 cortices from *Dhcr7^+/+^* or *Dhcr7^-/-^* embryos were dissected and mechanically dissociated into a single cell suspension by fire-polished glass pipette as previously described (Capecchi and Pozner, 2015; Tomita et al., 2020). Cell density and viability were determined using trypan blue exclusion. Cells were seeded in triplicate at a clonal density of 10 cells/μl in 6 well (2 ml/well) ultra-low attachment culture plates (Coster) in serum-free medium supplemented with 20 ng/ml EFG (Sigma), 20 ng/ml FGF2 (Sigma), 2% B27 supplement (Invitrogen) and 2 μg/ml heparin (Sigma). Neurospheres were cultured for 6 days at 37 °C. To evaluate self-renewal potential, neurospheres were mechanically dissociated into single cell suspensions by fire-polished glass pipette, passed through a 45μm nylon screen cell strainer, and cultured at a clonal density of 2 cells/μl for an additional 6 days.

### Immunocytochemistry and histological analysis

For morphometric analysis, immunostaining of tissue sections was performed as described (Capecchi and Pozner, 2015). Briefly, brain sections were permeabilized and blocked in PBST (1X PBS, 0.5% (v/v) Triton X-100) containing 10% NGS for 1 hour. Brain slices were incubated with primary antibodies in PBST with 5% NGS at 4 °C overnight. The sections were incubated with secondary antibodies in PBST with 5% NGS for 1-2 hours at room temperature. Sections were counterstained with DAPI (Thermo Fisher Scientific). Slides were mounted in Fluoromount-G anti-fade reagent (Southern Biotech). Digital image acquisition was performed with EVO-FL Imagining System (Thermo Fisher Scientific). For quantification of precursor and neuron numbers, we analyzed sections at the medial-lateral level, counting all marker-positive cells in a 200 µm wide strip of the cortex, extending from the meninges to the ventricle. In all cases, we analyzed at least 3 similar cortical sections/embryo or pup from 3 different embryos or pup per genotype (for a total of at least 9 sections per genotype).

### Antibodies

The primary antibodies used for immunostaining were chicken anti-GFP (1:1000; Abcam), rabbit anti-Dhcr7 (1:100; Thermo Fisher Scientific), rabbit anti-Sox2 (1:200; Cell Signaling Technology), rabbit anti-Pax6 (1:1000; Covance), mouse anti-Ki67 (1:200; BD Biosciences), mouse anti-βIII-tubulin (1:1000; Covance), rabbit anti-βIII-tubulin (1:1000; Covance), rabbit anti-Tbr1 (1:500; Abcam), mouse anti-Satb2 (1:400; Abcam), rat anti-Ctip2 (1:500; Abcam), and rabbit anti-cleaved caspase 3 (1:400; Cell Signaling Technology), mouse anti-Nestin (1:1000; StemCell Technologies), rabbit anti-Brachyury (1:200; R&D Systems), goat anti-NCAM (1:200; R&D Systems), goat anti-Sox17 (1:200; R&D Systems), rabbit anti-Foxa2 (1:1000; Abcam). The secondary antibodies used for immunostaining were Rhodamine (TRITC)-conjugated goat anti-mouse and anti-rabbit IgG (1:500; Jackson ImmunoResearch Laboratories) and Alexa Fluor 488-conjugated goat anti-mouse, anti-rat and anti-rabbit IgG (1:800; Jackson ImmunoResearch Laboratories), Alexa Flour 488-conjugated donkey anti-goat IgG (1:800; Jackson ImmunoResearch Laboratories), Rhodamine (TRITC)-conjugated donkey anti-goat (1:200; Jaclson ImmunoResearch Laboratories). The primary antibodies used for immunoblotting were rabbit anti-Dhcr7 (1:1000; Abcam), rabbit anti-GAPDH (1:5000; Santa Cruz Biotechnology), rabbit anti Erk1/2 (1:5000; Santa Cruz Biotechnology), rabbit anti-TrkB (1:1000; Cell Signaling Technology), rabbit anti-phospho-TrkB (1:1000; Cell Signaling Technology), rabbit anti-glucocorticoid receptor (1:1000; Cell Signaling Technology), rabbit anti-phospho-glucocorticoid receptor (1:1000; Cell Signaling Technology), rabbit anti-MEK (1:1000; Cell Signaling Technology), rabbit anti-phospho-MEK (1:1000; Cell Signaling), rabbit anti-phospho-cebpβ (1:1000; Cell signaling Technology) and rabbit anti-cebpβ (1:1000; Cell Signaling Technology), rabbit anti-Pax6 (1:1000; Covance), mouse anti-Nestin (1:1000; StemCell Technologies), rabbit anti-Brachyury (1:200; R&D Systems), rabbit anti-Tbr2 (1:500; Abcam), goat anti-Sox17 (1:200; R&D Systems), rabbit anti-Foxa2 (1:1000; Abcam). The secondary antibodies used for immunoblotting were HRP-conjugated goat anti-mouse IgG (1:5000; Jackson ImmunoResearch Laboratories) and anti-rabbit IgG (1:10,000; Jackson ImmunoResearch Laboratories), HRP-conjugated donkey anti-goat IgG (1:1000; Jackson ImmunoResearch Laboratories).

### RT-PCR

Total RNA was isolated with Trizol and cDNA was prepared using the SuperScript III Reverse Transcriptase kit (Invitrogen) according to the manufacture’s protocols. Primer sequences are the following: *Dhcr7*-F: 5’-TATGAGGTGAATGGGCTGCA-3’; *Dhcr7*-R: 5’-GGTTAATGAGGGTCCAGGCT -3’; *β-actin*-F: 5’-GATGACGATATCGCTGCGCTG-3’; *β-actin-*R: 5’-GTACGACCAGAGGCATACAGG-3’. All PCR products were single bands with predicted molecular weights and confirmed by DNA sequencing.

### Quantitative PCR

Total RNA was extracted with Tri-Reagent (Sigma) treated with DNAse I (Fermentas, Thermo Scientific, Waltham, MA, USA) and cDNA was synthesized from 1μg of RNA using the SuperScript IV Reverse Transcriptase Kit (Invitrogen) according to the manufacturer’s protocols. Quantitative PCR was performed using Taqman Fast Advance Mater Mix (Thermo Fisher Scientific) and Taqman probes targeted against either *Dhcr7* (Mm01164321_m1) or *β-Actin* (Mm00607939_s1). *β-Actin* mRNA was used as an endogenous control for all reactions, and all reactions were performed in triplicate. Quantitative PCR was performed and analyzed using StepOne Plus Real-Time PCR system (ThermoFisher Scientific).

### Western Blotting

Embryonic cortices or neurosphere cultures were lysed in RIPA buffer (50 mM Tri pH8, 150mM NaCl, 1% NP-40, 0.1% SDS, 1mM EDTA) containing 1mM PMSF (phenylmethanesulfonyl fluoride), 1mM sodium vanadate, 20mM sodium fluoride 10 μg/ml aprotinin and 10 μg/ml leupeptin. 10-20 μg of protein lysate was electrophoresed, and western blots were performed as described previously (Barnabe-Heider and Miller, 2003).

### RNA sequencing and data analysis

Raw RNA sequencing reads in FASTQ format were mapped to the human genome using HISAT (https://ccb.jhu.edu/software/hisat/; Last accessed January 22, 2021), and format conversions were performed using Samtools. Cufflinks (http://cole-trapnell-lab.github.io/cufflinks/; Last accessed January 22, 2021) was used to estimate relative abundances of transcripts from each RNA sample. Cuffdiff, a module of Cufflinks, was then used to determine differentially expressed genes (DEGs) between WT and KO hNPCs. DEGs met the following criteria: adjusted *p* value < .05 (corresponding to the allowed false discovery rate of 5%) and fold-change > 1.5 between genotypes. A two-way hierarchical clustering dendrogram (complete-linkage method, Euclidean distance scale) of DEGs was used to visualize biological variability among samples, generated by R software using the “pheatmap” package (https://cran.r-project.org/web/packages/pheatmap/). To elucidate biological functions of DEGs, the Core Analysis feature of Ingenuity Pathway Analysis (IPA®, Qiagen) was used to identify significantly enriched Diseases and Biological Functions related to the nervous system. Network interactions among DEGs involved in the Biological Function “development of the central nervous system” were assessed using STRING (Search Tool for the Retrieval of Interacting Genes/Proteins) analysis, set at the highest confidence interaction score and only connected nodes displayed (https://string-db.org/cgi/input.pl?sessionId=xCahIfrzvltC; Last accessed January 22, 2021). Enriched KEGG (Kyoto Encyclopedia of Genes and Genomes) pathways were identified among DEGs in the STRING network. Finally, a Venn diagram was generated to demonstrate the overlap between genes dysregulated in KO hNPCs and genes in the SFARI database, a collection of genes implicated in autism susceptibility (https://gene.sfari.org/; Last accessed January 22, 2021). Raw data of the RNA sequencing has been deposited at Dryad (https://doi.org/10.5061/dryad.zw3r2287f<u>; available upon publication</u>). The list of differentially expressed genes can be found in Supplementary File 2 (Excel).

### Oxysterol and Sterol Analysis

Cell pellets were resuspended in 300 μL of 1X PBS and lysed by sonication in an ice bath for 30 min. Protein determination was performed using the BioRad DC protein mass assay (BioRad, Hercules, CA). Internal standard mixtures for sterols and oxysterols analysis were added to each sample (see Tables S1-4 for a list of standards and their concentrations used). Lipid extraction was performed using the Folch method as described previously (Fliesler et al., 2018; Folch et al., 1957; Herron et al., 2018). The dried lipid extract was reconstituted with 200 μL of methylene chloride and stored at −80 °C until analysis. Prior to analysis, 50 μL of extract was transferred to into glass LC vials, dried under argon, and reconstituted with 50 μL of 90% methanol in water with 0.1% formic acid. For tissue samples, sterol and oxysterol internal standard mixtures were added to whole tissues, which were subsequently homogenized in 4 mL Folch solution with 1 mL 0.9% NaCl. The dried lipid extract was reconstituted with 0.5 mL (for tissues <50 mg), 1.0 mL (tissues >50 <100 mg) or 1.5 mL (>100 mg) of methylene chloride. Prior to analysis, 50 μL of lipid extract (30 μL for tissues >100 mg) was transferred into glass LC vials, dried under argon, and reconstituted with 50 μL 90% methanol in water with 0.1% formic acid. Determination of oxysterol and sterol concentrations in cells and tissues was performed by ultra-performance liquid chromatography (UPLC) tandem mass spectrometry (MS/MS) on a SCIEX 6500 triple quadrupole mass spectrometer (for oxysterols) or a SCIEX 4000 QTRAP (for sterols) mass spectrometer with atmospheric pressure chemical ionization (APCI) coupled to a Waters Acquity UPLC system, as described previously (Fliesler et al., 2018; Herron et al., 2018). Briefly, sterols and oxysterols were separated by reversed phase chromatography on a C18 column (1.7 mm, 2.1 x 100 mm, Phenomenex Kinetex) using a 15 min isocratic gradient of 90% methanol with 0.1% formic acid at a flow of 0.4 mL/min. Selective reaction monitoring (SRM) was used to monitor the dehydration of the sterol and oxysterol [M+H]^+^ ions to generate [M+H-H_2_O]^+^ ions (See Table S1-4 for a list of retention times and MS/MS transitions for the standards used). The APCI parameters were as follows: nebulizer current, 3 mA; temperature, 350 °C; curtain gas, 20 psi; ion source gas, 55 psi. The MS conditions for SRM analysis were as follows: declustering potential, 80 V; entrance potential, 10 V; collision energy, 25 V; collision cell exit potential, 20 V. Data analysis was performed with Analyst (v. 1.6.2) Quantitation Wizard. Analyte concentrations in cells and tissues were determined relative to the internal standard levels and the relative response factor (RRF) of each analyte calculated from a mixture of sterol and oxysterol standards and internal standards. Concentrations were normalized to cell protein weight or tissue weight.

### Docking simulation

Docking simulation was performed with Autodock Vina 1.1.2. The model ligands and 3D models were built and generated by Openbabel 2.3.2. Crystal structure of glucococorticoid receptor and docking of cortisol to glucocorticoid receptor were from crystal structure 4p6w available from Protein Data Bank (http://www.rcsb.org/structure/4p6w).

### Statistical analysis

In general, three biological replicates were used for each experiment based on the strong phenotype observed in the *Dhcr7*-KO SLOS model and our previous publication showing the large differences in oxysterols and sterol levels in WT and KO brains at birth (Xu et al., 2011). For culturing of mouse cortical precursors or neurospheres from single embryos, one litter is considered a biological replicate. For hNPC culture, each separate preparation of hNPC from hiPSC is considered a biological replicate. For *in vivo* study, one litter is considered a biological replicate. Statistical analyses were performed using two-tailed Student’s *t*-test assuming unequal variance when comparing two different groups unless otherwise indicated in the text. For immunostaining of cell culture or tissue sections, at least three technical replicates per biological replicate were performed. To analyze the multi-group neuroanatomical studies, we used one-way ANOVA unless otherwise indicated in the text. Significant interactions or main effects were further analyzed using Newman-Keuls post-hoc tests. All statistical tests were performed using Microsoft Excel or Prism 8 (GraphPad). In all cases, error bars indicate standard error of the mean.

## Figure Supplements

**Figure 1-Figure Supplement 1. Characterization of the pluripotency of SLOS-derived human iPSCs.** Related to Figure 1. Pluripotency of SLOS patient derived human iPSC lines, 3044 and 5788 were characterized. (A) Representative images of established SLOS patient derived human iPSC lines (3044 and 5788) and human iPSC line from healthy individual (emhf2). (B) RT-PCR analysis of hES cell marker genes in human iPSCs derived from SLOS patients (3044 and 5788) and healthy individual (emhf2). Primers used for Oct3/4, Sox2, Klf4 and c-Myc specifically amplified endogenous transcripts but not transgenic transcripts. (C) Immunostaining confirming in-vitro differentiation of SLOS patient derived human iPSCs (3044 and 5788) into all three germ layers. (D) RT-PCR analyses and Western blot analyses of differentiation markers for the three germ layers.

**Figure 1-Figure Supplement 2. Loss of *DHCR7* alleles causes decreased proliferation and increased neurogenesis in human cortical precursors.** (A) Images of cultured human cortical precursors immunostained for Dhcr7 (red) and Sox2 or Nestin (green). (B-D) Human SLOS-patient-derived (SLOS 3044 and SLOS 5788) and unaffected-patient-derived cortical precursors were cultured for 3 days and analyzed. (B) Cells were immunostained for Ki67 (green) and βIII-tubulin (red) after 3 days and quantified the proportions of Ki67+ (C) and βIII-tubulin+ cells (D). Scale Bar = 50 μm. *, *p* < 0.001; n = 3 biological replicates per genotype.

**Figure 2-Figure Supplement 1. Cholesterol precursor 7-DHC and 7-DHC-derived Oxysterols are accumulated *in* SLOS-derived human iPSCs and NPCs.** Related to Figure 2. SLOS patient derived hiPSCs and NPCs (3044 and 5788) along hiPSCs and NPC from healthy individual (emhf2) were cultured and used for LC-MS/MS analysis sterols and oxysterols. Quantitation of cholesterol, its precursors and accumulated 7-DHC derived oxysterols in SLOS patient derived hiPSCs and hiPSCs from healthy individuals (A) and in SLOS patient derived NPCs and NPCs from healthy individuals (B).

**Figure 3-Figure Supplement 1. Rescue of the neurogenesis phenotype in *Dhcr7*-knockdown mouse cortical precursors by human *DHCR7* cDNA expression vector.** Related to Figure 3. Potential off-target effects of murine *Dhcr7* shRNAs were determined. Human *DHCR7* cDNA expressing vectors transfected murine NPCs with either murine Dhcr7 shRNA vector or control shRNA vectors. The transfected cells were cultured for 3 days, immunostained and analyzed for proliferation marker, Ki67 or neuronal marker, βIII tubulin. (A) Cultures were immunostained for EGFP-mDhcr7shRNA and proliferation marker Ki67, the proportion of total EGFP+ cells that were also positive for Ki67 was quantified. (B) Cultures were immunostained for EGFP-mDhcr7shRNA and neuronal marker, βIII tubulin, the proportion of total EGFP+ cells that were also positive for βIII tubulin was quantified.

**Figure 7-Figure Supplement 1. Antioxidants rescue the neurogenic phenotype in human and murine NPCs with *Dhcr7* mutations.** Related to Figure 7. (A) Structures of DHCEO and known ligands of GR, OCDO and cortisol. (B, D) Another SLOS hNPC line (SLOS 5788) and mNPCs from E12.5 *Dhcr7*^-/-^ and *Dhcr7*^+/+^ embryonic cortices were cultured and treated with cholesterol, vitamin E, vitamin C, vitamin E/C or glutathione (GSH). The cells were immunostained for βIII-tubulin and DAPI three days after plated. (C) Quantification of the proportion of βIII-tubulin positive cells in wild type hNPCs and SLOS hNPCs treated with cholesterol or antioxidants. (D) Quantification of the proportion of βIII-tubulin positive cells in *Dhcr7*^-/-^ and *Dhcr7*^+/+^ NPCs treated with cholesterol or antioxidants. Error bars indicate SEM. *, *p*< 0.001. n = 3 per experiment. Scale Bar = 50 μm.

## Supplementary File 1

**Table S1. Retention times and MS/MS transitions for oxysterol internal standards. Relate to** Figure 2 **and S3.**

**Table S2. Retention times and MS/MS transitions for all oxysterol standards. Relate to** Figure 2 **and S3.**

**Table S3. Retention times and MS/MS transitions for sterol internal standards. Relate to Figure 2 and S3.**

**Table S4. Retention times and MS/MS transitions for sterol standards. Relate to** Figure 2 **and S3.**

**Table S5. Ingenuity Pathway Analysis (IPA®) reveals “development of the central nervous system” as one of the top 10 enriched Diseases and Biological Functions related to the nervous system.** Related to Figure S1. DEGs were further analyzed with (IPA®) to identify the most enriched biological functions related to the nervous system in SLOS mutant NPCs. The table below shows the top ten enriched terms corresponding to Diseases/Bio-functions along with the *p*-value and overlapping number of genes in the dataset.

**Figure S1. Perturbation in *DHCR7* causes gene expression changes in neurogenic pathways in human NPCs.** (A) Hierarchical clustering heatmap of differentially expressed genes show distinct expression pattern changes in transcript abundance for SLOS mutant NPCs as compared to wild type. Red color represents increase in abundance, blue color represents relative decrease, and white color represents no change. (B) Enriched KEGG pathways identified among DEGs from SLOS mutant NPCs and involved in the Biological Function “development of the central nervous system”, identified by Ingenuity Pathway Analysis (see Supporting Information). Color of bar corresponds to DEGs in the STRING network that are in enriched pathways. (C) String analysis of DEGs from SLOS mutant NPCs and involved in the Biological Function “Development of the central nervous system” identified by Ingenuity Pathway Analysis. Parameters of high confidence have been applied and only connected nodes are displayed. Arrows indicate whether the gene was up- or down-regulated. Color of DEG corresponds to enriched KEGG pathway. *n* =3 biological replicates per genotype; DEGs met the criteria of fold-change > 1.5 between genotypes and adjusted *p*-value < 0.05.

## Supplementary File 2

**Table S6 (Excel file). List of differentially expressed genes in SLOS hNPCs relative to wild-type hNPCs. Relate to Figure S1.**

## Notes

### Competing Interest Statement

The authors have declared no competing interest.

### Summary of Updates

Corrected a few typos in the main text and added a few sentences. Supplemental files updated.

## References

Barnabe-Heider, F., and Miller, F.D. (2003). Endogenously produced neurotrophins regulate survival and differentiation of cortical progenitors via distinct signaling pathways. J Neurosci 23, 5149–5160.

Bartkowska, K., Paquin, A., Gauthier, A.S., Kaplan, D.R., and Miller, F.D. (2007). Trk signaling regulates neural precursor cell proliferation and differentiation during cortical development. Development 134, 4369–4380. doi: 10.1242/dev.008227.

Bonni, A., Brunet, A., West, A.E., Datta, S.R., Takasu, M.A., and Greenberg, M.E. (1999). Cell survival promoted by the Ras-MAPK signaling pathway by transcription-dependent and - independent mechanisms. Science 286, 1358–1362. doi: 10.1126/science.286.5443.1358.

Bukelis, I., Porter, F.D., Zimmerman, A.W., and Tierney, E. (2007). Smith-Lemli-Opitz syndrome and autism spectrum disorder. Am J Psychiatry 164, 1655–1661. doi: 164/11/1655 [pii] 10.1176/appi.ajp.2007.07020315.

Byrne, E.F., Sircar, R., Miller, P.S., Hedger, G., Luchetti, G., Nachtergaele, S., Tully, M.D., Mydock-McGrane, L., Covey, D.F., Rambo, R.P., et al. (2016). Structural basis of Smoothened regulation by its extracellular domains. Nature 535, 517–522. doi: 10.1038/nature18934.

Capecchi, M.R., and Pozner, A. (2015). ASPM regulates symmetric stem cell division by tuning Cyclin E ubiquitination. Nat Commun 6, 8763. doi: 10.1038/ncomms9763.

Chenn, A., and Walsh, C.A. (2002). Regulation of cerebral cortical size by control of cell cycle exit in neural precursors. Science 297, 365–369. doi: 10.1126/science.1074192.

Corcoran, R.B., and Scott, M.P. (2006). Oxysterols stimulate Sonic hedgehog signal transduction and proliferation of medulloblastoma cells. Proc Natl Acad Sci U S A 103, 8408–8413. doi: 10.1073/pnas.0602852103.

Dietschy, J.M., and Turley, S.D. (2004). Cholesterol metabolism in the central nervous system during early development and in the mature animal. J Lipid Res 45, 1375–1397. doi: 10.1194/jlr.R400004-JLR200 R400004-JLR200 [pii].

DiSalvo, C.V., Zhang, D., and Jacobberger, J.W. (1995). Regulation of NIH-3T3 cell G1 phase transit by serum during exponential growth. Cell Prolif 28, 511–524. doi: 10.1111/j.1365-2184.1995.tb00089.x.

Doba, T., Burton, G.W., and Ingold, K.U. (1985). Antioxidant and co-antioxidant activity of vitamin C. The effect of vitamin C, either alone or in the presence of vitamin E or a water-soluble vitamin E analogue, upon the peroxidation of aqueous multilamellar phospholipid liposomes. Biochimica et Biophysica Acta (BBA) - Lipids and Lipid Metabolism 835, 298–303. doi: https://doi.org/10.1016/0005-2760(85)90285-1.

DuSell, C.D., Umetani, M., Shaul, P.W., Mangelsdorf, D.J., and McDonnell, D.P. (2008). 27-Hydroxycholesterol Is an Endogenous Selective Estrogen Receptor Modulator. Molecular Endocrinology 22, 65–77. doi: 10.1210/me.2007-0383.

Ferreira, L.G., Dos Santos, R.N., Oliva, G., and Andricopulo, A.D. (2015). Molecular docking and structure-based drug design strategies. Molecules 20, 13384–13421. doi: 10.3390/molecules200713384.

Fitzky, B.U., Moebius, F.F., Asaoka, H., Waage-Baudet, H., Xu, L., Xu, G., Maeda, N., Kluckman, K., Hiller, S., Yu, H., et al. (2001). 7-Dehydrocholesterol-dependent proteolysis of HMG-CoA reductase suppresses sterol biosynthesis in a mouse model of Smith-Lemli-Opitz/RSH syndrome. J Clin Invest 108, 905–915. doi: 10.1172/jci12103.

Fitzky, B.U., Witsch-Baumgartner, M., Erdel, M., Lee, J.N., Paik, Y.K., Glossmann, H., Utermann, G., and Moebius, F.F. (1998). Mutations in the Delta 7-sterol reductase gene in patients with the Smith-Lemli-Opitz syndrome. Proc Natl Acad Sci U S A 95, 8181–8186.

Fliesler, S.J., Peachey, N.S., Herron, J., Hines, K.M., Weinstock, N.I., Ramachandra Rao, S., and Xu, L. (2018). Prevention of Retinal Degeneration in a Rat Model of Smith-Lemli-Opitz Syndrome. Sci Rep 8, 1286. doi: 10.1038/s41598-018-19592-8.

Folch, J., Lees, M., and Sloane Stanley, G.H. (1957). A simple method for the isolation and purification of total lipides from animal tissues. J Biol Chem 226, 497–509.

Francis, K.R., Ton, A.N., Xin, Y., O’Halloran, P.E., Wassif, C.A., Malik, N., Williams, I.M., Cluzeau, C.V., Trivedi, N.S., Pavan, W.J., et al. (2016). Modeling Smith-Lemli-Opitz syndrome with induced pluripotent stem cells reveals a causal role for Wnt/beta-catenin defects in neuronal cholesterol synthesis phenotypes. Nat Med 22, 388–396. doi: 10.1038/nm.4067.

Herron, J., Hines, K.M., and Xu, L. (2018). Assessment of Altered Cholesterol Homeostasis by Xenobiotics Using Ultra-High Performance Liquid Chromatography-Tandem Mass Spectrometry. Curr Protoc Toxicol 78, e65. doi: 10.1002/cptx.65.

Huang, P., Zheng, S., Wierbowski, B.M., Kim, Y., Nedelcu, D., Aravena, L., Liu, J., Kruse, A.C., and Salic, A. (2018). Structural Basis of Smoothened Activation in Hedgehog Signaling. Cell 174, 312–324. doi: 10.1016/j.cell.2018.04.029.

Jeanneteau, F., Garabedian, M.J., and Chao, M.V. (2008). Activation of Trk neurotrophin receptors by glucocorticoids provides a neuroprotective effect. Proc Natl Acad Sci U S A 105, 4862–4867. doi: 10.1073/pnas.0709102105.

Jiang, X.S., Wassif, C.A., Backlund, P.S., Song, L., Holtzclaw, L.A., Li, Z., Yergey, A.L., and Porter, F.D. (2010). Activation of Rho GTPases in Smith-Lemli-Opitz syndrome: pathophysiological and clinical implications. Hum Mol Genet 19, 1347–1357. doi: 10.1093/hmg/ddq011.

Konopka, G., Wexler, E., Rosen, E., Mukamel, Z., Osborn, G.E., Chen, L., Lu, D., Gao, F., Gao, K., Lowe, J.K., et al. (2012). Modeling the functional genomics of autism using human neurons. Molecular Psychiatry 17, 202–214. doi: 10.1038/mp.2011.60.

Korade, Z., Xu, L., Harrison, F.E., Ahsen, R., Hart, S.E., Folkes, O.M., Mirnics, K., and Porter, N.A. (2014). Antioxidant Supplementation Ameliorates Molecular Deficits in Smith-Lemli-Opitz Syndrome. Biol Psychiatry 75, 215–222. doi: S0006-3223(13)00591-X [pii] 10.1016/j.biopsych.2013.06.013.

Korade, Z., Xu, L., Shelton, R., and Porter, N.A. (2010). Biological activities of 7-dehydrocholesterol-derived oxysterols: implications for Smith-Lemli-Opitz syndrome. J Lipid Res 51, 3259–3269. doi: jlr.M009365 [pii] 10.1194/jlr.M009365.

Menard, C., Hein, P., Paquin, A., Savelson, A., Yang, X.M., Lederfein, D., Barnabe-Heider, F., Mir, A.A., Sterneck, E., Peterson, A.C., et al. (2002). An essential role for a MEK-C/EBP pathway during growth factor-regulated cortical neurogenesis. Neuron 36, 597–610. doi: 10.1016/s0896-6273(02)01026-7.

Molyneaux, B.J., Arlotta, P., Menezes, J.R., and Macklis, J.D. (2007). Neuronal subtype specification in the cerebral cortex. Nat Rev Neurosci 8, 427–437. doi: 10.1038/nrn2151.

Myers, B.R., Neahring, L., Zhang, Y., Roberts, K.J., and Beachy, P.A. (2017). Rapid, direct activity assays for Smoothened reveal Hedgehog pathway regulation by membrane cholesterol and extracellular sodium. Proc Natl Acad Sci U S A 114, E11141–E11150. doi: 10.1073/pnas.1717891115.

Okita, K., Matsumura, Y., Sato, Y., Okada, A., Morizane, A., Okamoto, S., Hong, H., Nakagawa, M., Tanabe, K., Tezuka, K., et al. (2011). A more efficient method to generate integration-free human iPS cells. Nat Methods 8, 409–412. doi: 10.1038/nmeth.1591.

Olkku, A., and Mahonen, A. (2009). Calreticulin mediated glucocorticoid receptor export is involved in β-catenin translocation and Wnt signalling inhibition in human osteoblastic cells. Bone 44, 555–565. doi: https://doi.org/10.1016/j.bone.2008.11.013.

Porter, F.D., and Herman, G.E. (2011). Malformation syndromes caused by disorders of cholesterol synthesis. J Lipid Res 52, 6–34. doi: jlr.R009548 [pii] 10.1194/jlr.R009548.

Porter, J.A., Young, K.E., and Beachy, P.A. (1996). Cholesterol modification of hedgehog signaling proteins in animal development. Science 274, 255–259.

Porter, N.A., Xu, L., and Pratt, D.A. (2020). Reactive Sterol Electrophiles: Mechanisms of Formation and Reactions with Proteins and Amino Acid Nucleophiles. Chemistry 2, 390–417. doi: 10.3390/chemistry2020025.

Raleigh, D.R., Sever, N., Choksi, P.K., Sigg, M.A., Hines, K.M., Thompson, B.M., Elnatan, D., Jaishankar, P., Bisignano, P., Garcia-Gonzalo, F.R., et al. (2018). Cilia-Associated Oxysterols Activate Smoothened. Mol Cell 72, 316–327 e315. doi: 10.1016/j.molcel.2018.08.034.

Sever, N., Mann, R.K., Xu, L., Snell, W.J., Hernandez-Lara, C.I., Porter, N.A., and Beachy, P.A. (2016). Endogenous B-ring oxysterols inhibit the Hedgehog component Smoothened in a manner distinct from cyclopamine or side-chain oxysterols. Proc Natl Acad Sci U S A 113, 5904–5909. doi: 10.1073/pnas.1604984113.

Sheng, R., Kim, H., Lee, H., Xin, Y., Chen, Y., Tian, W., Cui, Y., Choi, J.C., Doh, J., Han, J.K., et al. (2014). Cholesterol selectively activates canonical Wnt signalling over non-canonical Wnt signalling. Nat Commun 5, 4393. doi: 10.1038/ncomms5393.

Sikora, D., Pettit-Kekel, K., Penfield, J., Merkens, L., and Steiner, R. (2006). The near universal presence of autism spectrum disorders in children with Smith-Lemli-Opitz syndrome. Am J Med Genet 140A, 1511-1518. doi: 10.1002/(issn)1552-4833.

Theofilopoulos, S., Wang, Y., Kitambi, S.S., Sacchetti, P., Sousa, K.M., Bodin, K., Kirk, J., Salto, C., Gustafsson, M., Toledo, E.M., et al. (2013). Brain endogenous liver X receptor ligands selectively promote midbrain neurogenesis. Nat Chem Biol 9, 126–133. doi: 10.1038/nchembio.1156.

Thurm, A., Tierney, E., Farmer, C., Albert, P., Joseph, L., Swedo, S., Bianconi, S., Bukelis, I., Wheeler, C., Sarphare, G., et al. (2016). Development, behavior, and biomarker characterization of Smith-Lemli-Opitz syndrome: an update. J Neurodev Disord 8, 12. doi: 10.1186/s11689-016-9145-x.

Tierney, E., Bukelis, I., Thompson, R.E., Ahmed, K., Aneja, A., Kratz, L., and Kelley, R.I. (2006). Abnormalities of cholesterol metabolism in autism spectrum disorders. Am J Med Genet B 141B, 666-668. doi: 10.1002/ajmg.b.30368.

Tint, G.S., Irons, M., Elias, E.R., Batta, A.K., Frieden, R., Chen, T.S., and Salen, G. (1994). Defective cholesterol biosynthesis associated with the Smith-Lemli-Opitz syndrome. N Engl J Med 330, 107–113.

Tint, G.S., Seller, M., Hughes-Benzie, R., Batta, A.K., Shefer, S., Genest, D., Irons, M., Elias, E., and Salen, G. (1995). Markedly increased tissue concentrations of 7-dehydrocholesterol combined with low levels of cholesterol are characteristic of the Smith-Lemli-Opitz syndrome. J Lipid Res 36, 89–95.

Tomita, H., Cornejo, F., Aranda-Pino, B., Woodard, C.L., Rioseco, C.C., Neel, B.G., Alvarez, A.R., Kaplan, D.R., Miller, F.D., and Cancino, G.I. (2020). The Protein Tyrosine Phosphatase Receptor Delta Regulates Developmental Neurogenesis. Cell Rep 30, 215–228 e215. doi: 10.1016/j.celrep.2019.11.033.

Usui, N., Araujo, D.J., Kulkarni, A., Co, M., Ellegood, J., Harper, M., Toriumi, K., Lerch, J.P., and Konopka, G. (2017). Foxp1 regulation of neonatal vocalizations via cortical development. Genes Dev 31, 2039–2055. doi: 10.1101/gad.305037.117.

Voisin, M., de Medina, P., Mallinger, A., Dalenc, F., Huc-Claustre, E., Leignadier, J., Serhan, N., Soules, R., Ségala, G., Mougel, A., et al. (2017). Identification of a tumor-promoter cholesterol metabolite in human breast cancers acting through the glucocorticoid receptor. Proceedings of the National Academy of Sciences 114, E9346. doi: 10.1073/pnas.1707965114.

Wassif, C.A., Maslen, C., Kachilele-Linjewile, S., Lin, D., Linck, L.M., Connor, W.E., Steiner, R.D., and Porter, F.D. (1998). Mutations in the human sterol Delta(7)-reductase gene at 11q12-13 cause Smith-Lemli-Opitz syndrome. American Journal of Human Genetics 63, 55–62.

Xu, L., Davis, T.A., and Porter, N.A. (2009). Rate Constants for Peroxidation of Polyunsaturated Fatty Acids and Sterols in Solution and in Liposomes. J Am Chem Soc 131, 13037–13044.

Xu, L., Korade, Z., and Porter, N.A. (2010). Oxysterols from free radical chain oxidation of 7-dehydrocholesterol: product and mechanistic studies. J Am Chem Soc 132, 2222–2232. doi: 10.1021/ja9080265.

Xu, L., Korade, Z., Rosado, D.A., Jr., Mirnics, K., and Porter, N.A. (2013). Metabolism of oxysterols derived from nonenzymatic oxidation of 7-dehydrocholesterol in cells. J Lipid Res 54, 1135–1143. doi: 10.1194/jlr.M035733 jlr.M035733 [pii].

Xu, L., Korade, Z., Rosado, D.A., Liu, W., Lamberson, C.R., and Porter, N.A. (2011). An oxysterol biomarker for 7-dehydrocholesterol oxidation in cell/mouse models for Smith-Lemli-Opitz syndrome. J Lipid Res 52, 1222–1233. doi: jlr.M014498 [pii] 10.1194/jlr.M014498.

Xu, L., Mirnics, K., Bowman, A.B., Liu, W., Da, J., Porter, N.A., and Korade, Z. (2012). DHCEO accumulation is a critical mediator of pathophysiology in a Smith-Lemli-Opitz syndrome model. Neurobiol Dis 45, 923–929. doi: S0969-9961(11)00386-X [pii] 10.1016/j.nbd.2011.12.011.

Yang, S.H., Sharrocks, A.D., and Whitmarsh, A.J. (2013). MAP kinase signalling cascades and transcriptional regulation. Gene 513, 1–13. doi: 10.1016/j.gene.2012.10.033.

Yin, H., Xu, L., and Porter, N.A. (2011). Free radical lipid peroxidation: mechanisms and analysis. Chem Rev 111, 5944–5972. doi: 10.1021/cr200084z.

Yu, J., Vodyanik, M.A., Smuga-Otto, K., Antosiewicz-Bourget, J., Frane, J.L., Tian, S., Nie, J., Jonsdottir, G.A., Ruotti, V., Stewart, R., et al. (2007). Induced pluripotent stem cell lines derived from human somatic cells. Science 318, 1917–1920. doi: 10.1126/science.1151526.

Zhou, H., Mehta, S., Srivastava, S.P., Grabinska, K., Zhang, X., Wong, C., Hedayat, A., Perrotta, P., Fernandez-Hernando, C., Sessa, W.C., et al. (2020). Endothelial cell-glucocorticoid receptor interactions and regulation of Wnt signaling. JCI Insight 5. doi: 10.1172/jci.insight.131384.

