## Supplementary material for "7-Dehydrocholesterol-derived oxysterols cause neurogenic defects in Smith-Lemli-Opitz syndrome": Figure Supplements related to main figures

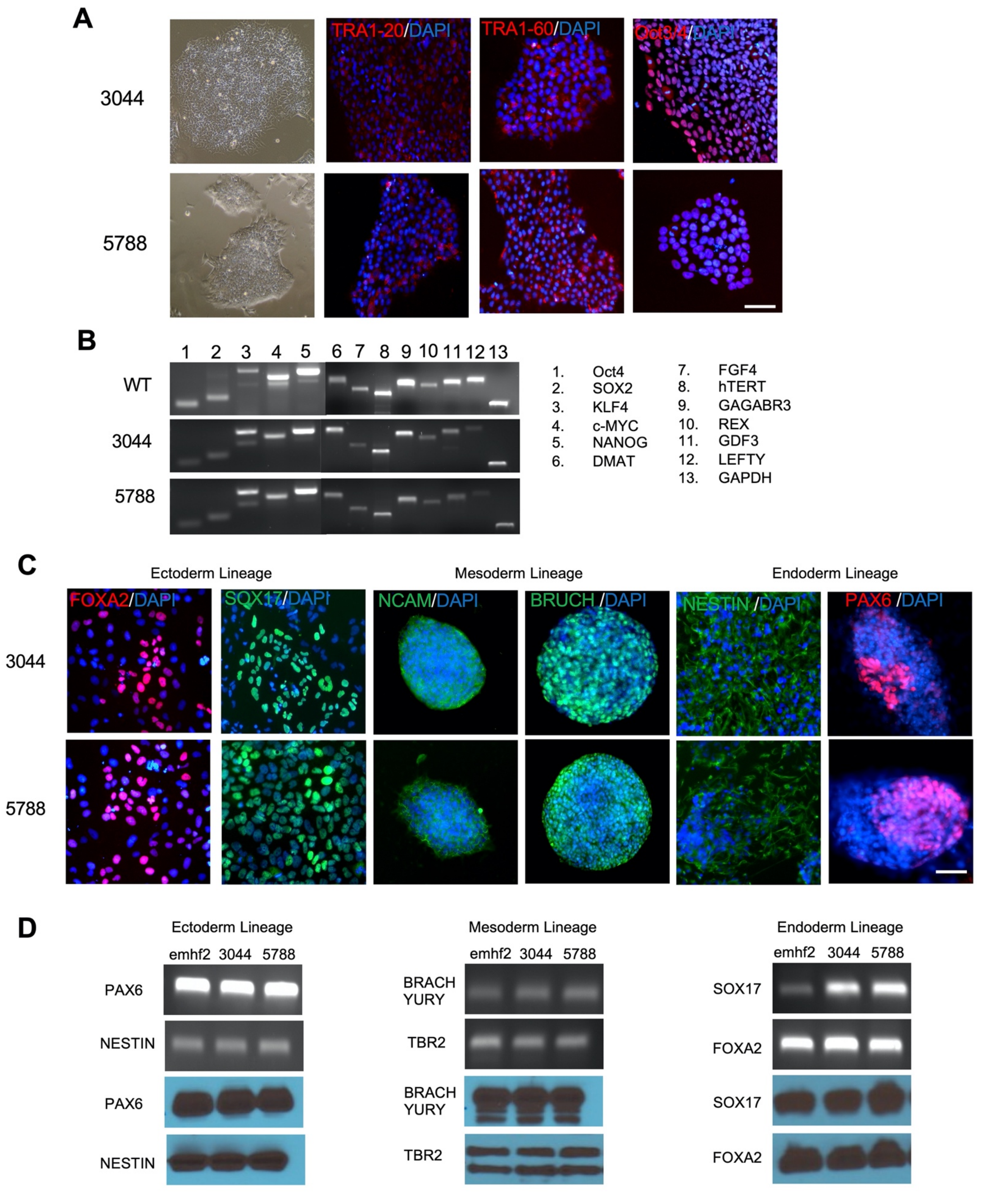


**Figure 1-Figure Supplement 1. Characterization of the pluripotency of SLOS-derived human iPSCs.** Related to Figure 1. Pluripotency of SLOS patient derived human iPSC lines, 3044 and 5788 were characterized. (A) Representative images of established SLOS patient derived human iPSC lines (3044 and 5788) and human iPSC line from healthy individual (emhf2). (B) RT-PCR analysis of hES cell marker genes in human iPSCs derived from SLOS patients (3044 and 5788) and healthy individual (emhf2). Primers used for Oct3/4, Sox2, Klf4 and c-Myc specifically amplified endogenous transcripts but not transgenic transcripts. (C) Immunostaining confirming in-vitro differentiation of SLOS patient derived human iPSCs (3044 and 5788) into all three germ layers. (D) RT-PCR analyses and Western blot analyses of differentiation markers for the three germ layers.

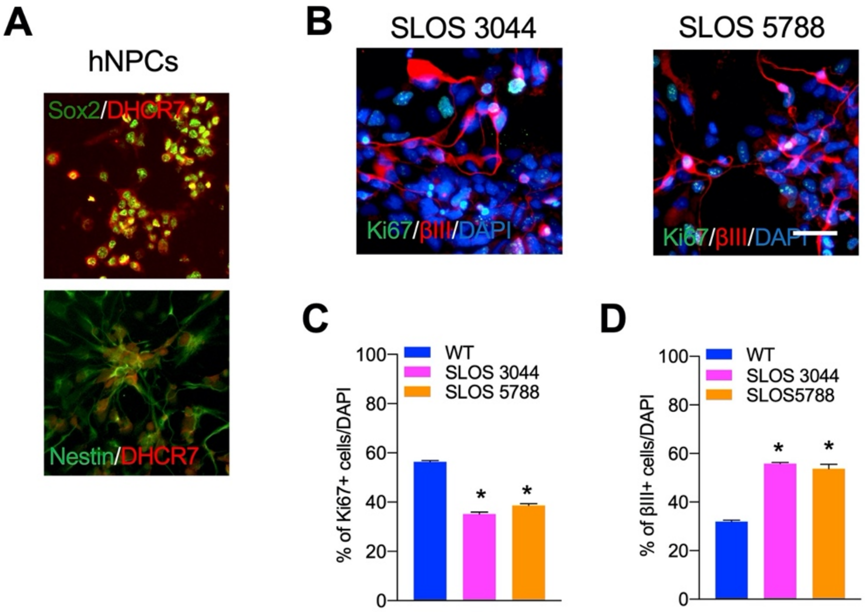


**Figure 1-Figure Supplement 2. Loss of *DHCR7* alleles causes decreased proliferation and increased neurogenesis in human cortical precursors.** (A) Images of cultured human cortical precursors immunostained for Dhcr7 (red) and Sox2 or Nestin (green). (B-D) Human SLOS-patient-derived (SLOS 3044 and SLOS 5788) and unaffected-patient-derived cortical precursors were cultured for 3 days and analyzed. (B) Cells were immunostained for Ki67 (green) and βIII-tubulin (red) after 3 days and quantified the proportions of Ki67+ (C) and βIII-tubulin+ cells (D). Scale Bar = 50 μm. *, $p < 0.001$; n = 3 biological replicates per genotype.

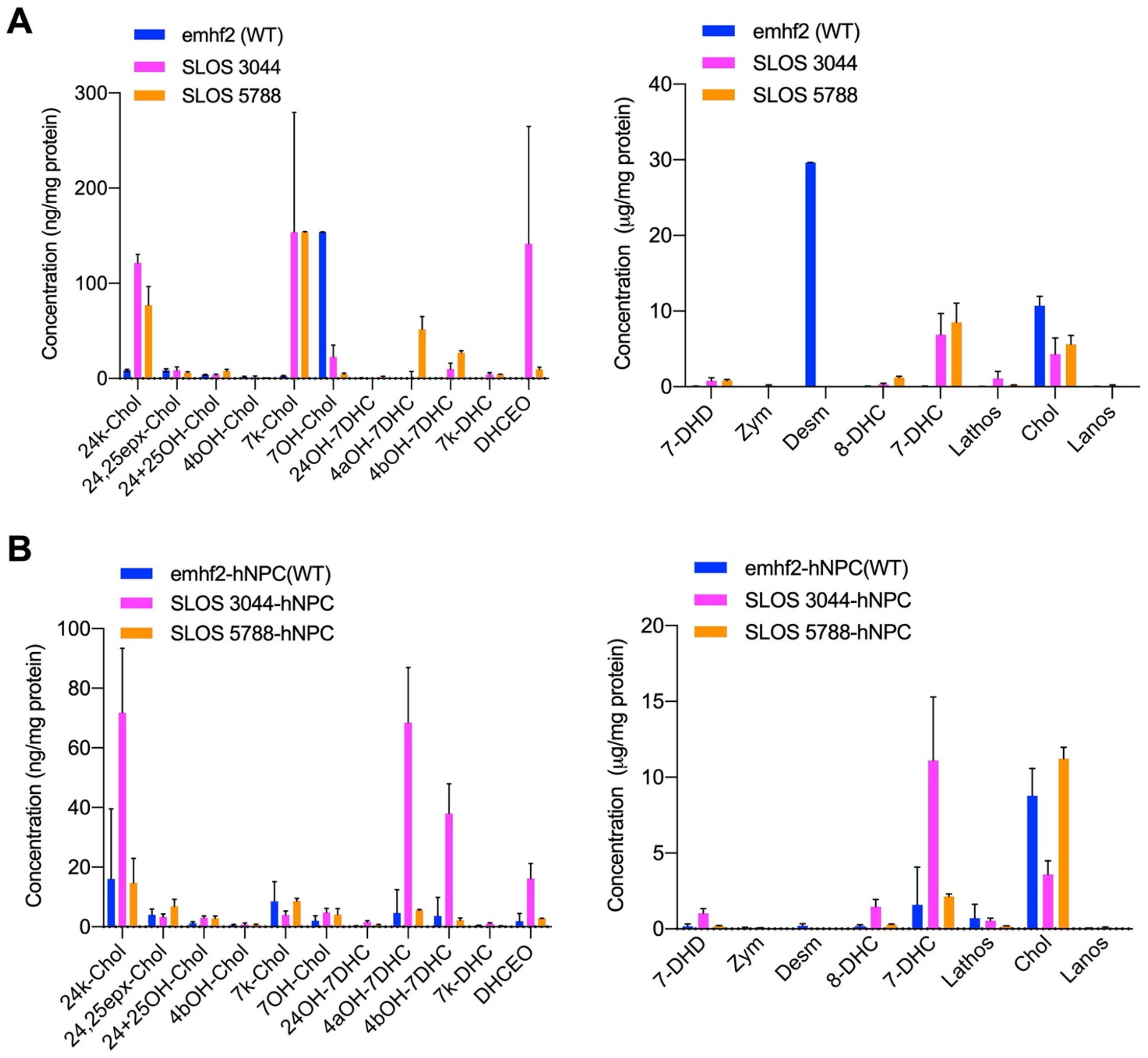


**Figure 2-Figure Supplement 1. Cholesterol precursor 7-DHC and 7-DHC-derived Oxysterols are accumulated *in* SLOS-derived human iPSCs and NPCs.** Related to Figure 2. SLOS patient derived hiPSCs and NPCs (3044 and 5788) along hiPSCs and NPC from healthy individual (emhf2) were cultured and used for LC-MS/MS analysis sterols and oxysterols. Quantitation of cholesterol, its precursors and accumulated 7-DHC derived oxysterols in SLOS patient derived hiPSCs and hiPSCs from healthy individuals (A) and in SLOS patient derived NPCs and NPCs from healthy individuals (B).

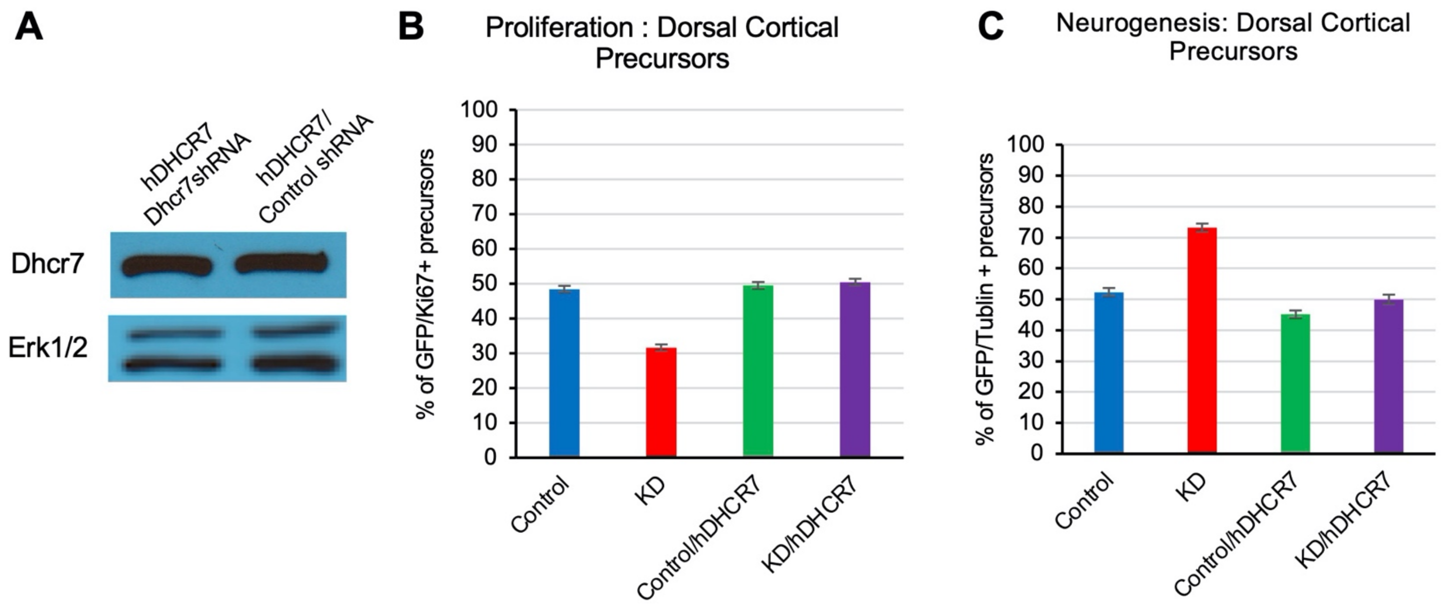


**Figure 3-Figure Supplement 1. Rescue of the neurogenesis phenotype in *Dhcr7*-knockdown mouse cortical precursors by human *DHCR7* cDNA expression vector.** Related to Figure 3. Potential off-target effects of murine *Dhcr7* shRNAs were determined. Human *DHCR7* cDNA expressing vectors transfected murine NPCs with either murine Dhcr7 shRNA vector or control shRNA vectors. The transfected cells were cultured for 3 days, immunostained and analyzed for proliferation marker, Ki67 or neuronal marker, βIII tubulin. (A) Cultures were immunostained for EGFP-mDhcr7shRNA and proliferation marker Ki67, the proportion of total EGFP+ cells that were also positive for Ki67 was quantified. (B) Cultures were immunostained for EGFP-mDhcr7shRNA and neuronal marker, βIII tubulin, the proportion of total EGFP+ cells that were also positive for βIII tubulin was quantified.

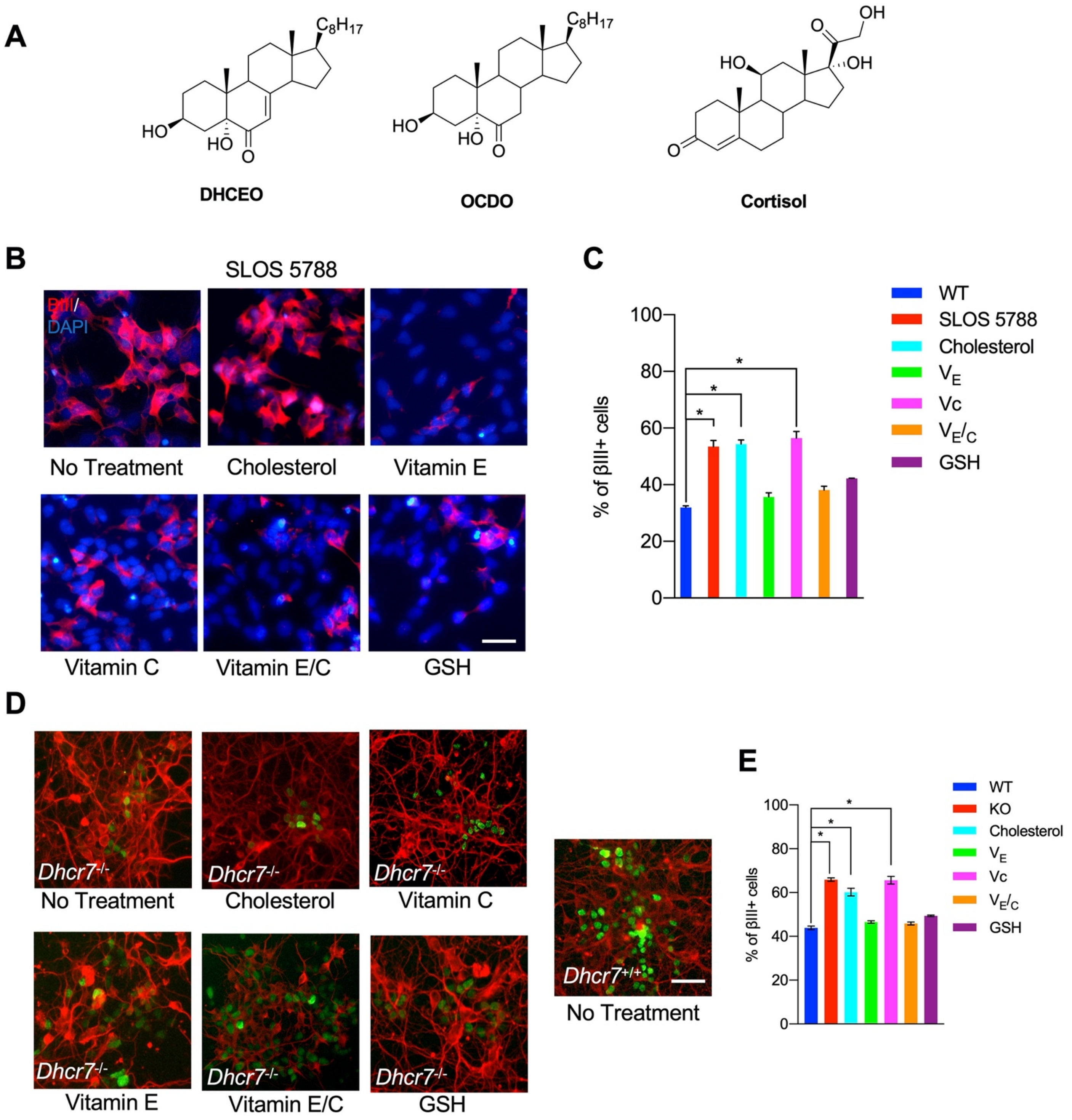


**Figure 7-Figure Supplement 1. Antioxidants rescue the neurogenic phenotype in human and murine NPCs with *Dhcr7* mutations.** Related to Figure 7. (A) Structures of DHCEO and known ligands of GR, OCDO and cortisol. (B, D) Another SLOS hNPC line (SLOS 5788) and mNPCs from E12.5 *Dhcr7*$^{-/-}$ and *Dhcr7*$^{+/+}$ embryonic cortices were cultured and treated with cholesterol, vitamin E, vitamin C, vitamin E/C or glutathione (GSH). The cells were immunostained for βIII-tubulin and DAPI three days after plated. (C) Quantification of the proportion of βIII-tubulin positive cells in wild type hNPCs and SLOS hNPCs treated with cholesterol or antioxidants. (D) Quantification of the proportion of βIII-tubulin positive cells in *Dhcr7*$^{-/-}$ and *Dhcr7*$^{+/+}$ NPCs treated with cholesterol or antioxidants. Error bars indicate SEM. *, $p< 0.001$. n = 3 per experiment. Scale Bar = 50 μm.
