## Supplemental Tables and Figures for "7-Dehydrocholesterol-derived oxysterols cause neurogenic defects in Smith-Lemli-Opitz syndrome"

**Table S1. Retention times and MS/MS transitions for oxysterol internal standards. Relate to Figure 2 and S3.**

| Internal Standard | Retention Time (min) | Q1 | Q3 |
|---|---|---|---|
| $d_7$-DHCEO | 1.80 | 424.3 | 406.3 |
| $d_7$-7-ketocholesterol | 2.91 | 408.3 | 390.3 |
| $d_6$-24,25-epoxycholesterol | 1.94 | 389.3 | 371.3 |
| $d_7$-24-hydroxycholesterol | 1.63 | 392.3 | 374.3 |
| $d_7$-4β-hydroxycholesterol | 5.16 | 392.3 | 374.3 |

**Table S2. Retention times and MS/MS transitions for all oxysterol standards. Relate to Figure 2 and S3.**

| Analyte | Retention Time (min) | Q1 | Q3 | Std Conc. Std+IS Mix | IS Used | IS Conc. Std+IS Mix |
|---|---|---|---|---|---|---|
| 24-hydroxycholesterol | 1.63 | 385.3 | 367.3 | 0.2 ug/mL | $d_7$-24-hydroxycholesterol | 0.2 ug/mL |
| 25-hydroxycholesterol | 1.63 | 385.3 | 367.3 | 0.2 ug/mL | $d_7$-24-hydroxycholesterol | 0.2 ug/mL |
| 24-ketocholesterol | 1.79 | 383.3 | 365.3 | 0.2 ug/mL | $d_6$-24,25-epoxycholesterol | 0.2 ug/mL |
| 24-epoxycholesterol | 1.94 | 383.3 | 365.3 | 0.2 ug/mL | $d_6$-24,25-epoxycholesterol | 0.2 ug/mL |
| 7β-hydroxycholesterol | 2.65 | 385.3 | 367.3 | 0.2 ug/mL | $d_7$-7-ketocholesterol | 0.2 ug/mL |
| 7α-hydroxycholesterol | 2.65 | 385.3 | 367.3 | 0.2 ug/mL | $d_7$-7-ketocholesterol | 0.2 ug/mL |
| 7-ketocholesterol | 2.96 | 401.3 | 383.3 | 0.2 ug/mL | $d_7$-7-ketocholesterol | 0.2 ug/mL |
| 4β-hydroxycholesterol | 5.16 | 385.3 | 367.3 | 0.2 ug/mL | $d_7$-4β-hydroxycholesterol | 0.2 ug/mL |
| 24-hydroxy-7-DHC | 1.43 | 383.3 | 365.3 | 0.2 ug/mL | $d_7$-DHCEO | 0.2 ug/mL |
| DHCEO | 1.83 | 417.3 | 399.3 | 0.2 ug/mL | $d_7$-DHCEO | 0.2 ug/mL |
| 7-keto-DHC | 2.14 | 399.3 | 381.3 | 0.2 ug/mL | $d_7$-DHCEO | 0.2 ug/mL |
| 4α-hydroxy-7-DHC | 3.77 | 365.3 | 365.3 | 0.2 ug/mL | $d_7$-DHCEO | 0.2 ug/mL |
| 4β-hydroxy-7-DHC | 4.37 | 365.3 | 365.3 | 0.2 ug/mL | $d_7$-DHCEO | 0.2 ug/mL |

**Table S3. Retention times and MS/MS transitions for sterol internal standards. Relate to Figure 2 and S3.**

| Internal Standard | Retention Time (min) | Q1 | Q3 |
|---|---|---|---|
| $^{13}C_3$-desmosterol | 6.40 | 370.3 | 370.3 |
| $d_7$-7-dehydrocholesterol | 6.90 | 374.3 | 374.3 |
| $d_7$-cholesterol | 8.60 | 376.3 | 376.3 |
| $^{13}C_3$-lanosterol | 9.83 | 412.3 | 412.3 |

**Table S4. Retention times and MS/MS transitions for sterol standards. Relate to Figure 2 and S3.**

| Analyte | Retention Time (min) | Q1 | Q3 | Std Conc. Std+IS Mix | IS Used | IS Conc. Std+IS Mix |
|---|---|---|---|---|---|---|
| 7-Dehydrodesmosterol | 5.18 | 365.3 | 365.3 | 0.4 ug/mL | $^{13}C_3$-desmosterol | 0.4 ug/mL |
| Zymosterol | 6.01 | 367.3 | 367.3 | 0.4 ug/mL | $^{13}C_3$-desmosterol | 0.4 ug/mL |
| Desmosterol | 6.38 | 367.3 | 367.3 | 0.4 ug/mL | $^{13}C_3$-desmosterol | 0.4 ug/mL |
| 8-Dehydrocholesterol | 6.79 | 367.3 | 367.3 | 0.4 ug/mL | $d_7$-7-dehydrocholesterol | 2.0 ug/mL |
| 7-Dehydrocholesterol | 7.0 | 367.3 | 367.3 | 0.4 ug/mL | $d_7$-7-dehydrocholesterol | 2.0 ug/mL |
| Lathosterol | 8.28 | 369.3 | 369.3 | 0.4 ug/mL | $^{13}C_3$-lanosterol | 0.4 ug/mL |
| Cholesterol | 8.69 | 369.3 | 369.3 | 0.4 ug/mL | $d_7$-cholesterol | 2.0 ug/mL |
| Lanosterol | 9.86 | 409.3 | 409.3 | 0.4 ug/mL | $^{13}C_3$-lanosterol | 0.4 ug/mL |

**Table S5. Ingenuity Pathway Analysis (IPA®) reveals "development of the central nervous system" as one of the top 10 enriched Diseases and Biological Functions related to the nervous system.** DEGs were further analyzed with (IPA®) to identify the most enriched biological functions related to the nervous system in SLOS mutant NPCs. The table below shows the top ten enriched terms corresponding to Diseases/Bio-functions along with the *p*-value and overlapping number of genes in the dataset.

| **Disease or Function** | **p-Value** | **# genes in dataset** |
|---|---|---|
| Development of neurons | 1.59E-47 | 275 |
| Morphology of nervous system | 4.71E-47 | 311 |
| Abnormal morphology of nervous system | 7.45E-42 | 265 |
| Development of sensory organ | 9.01E-33 | 186 |
| Development of central nervous system | 3.28E-29 | 199 |
| Neurotransmission | 6.97E-29 | 143 |
| Abnormal morphology of central nervous system | 1.42E-28 | 162 |
| Morphogenesis of neurons | 1.53E-28 | 192 |
| Neuritogenesis | 5.85E-28 | 189 |
| Morphology of central nervous system | 7.02E-28 | 184 |

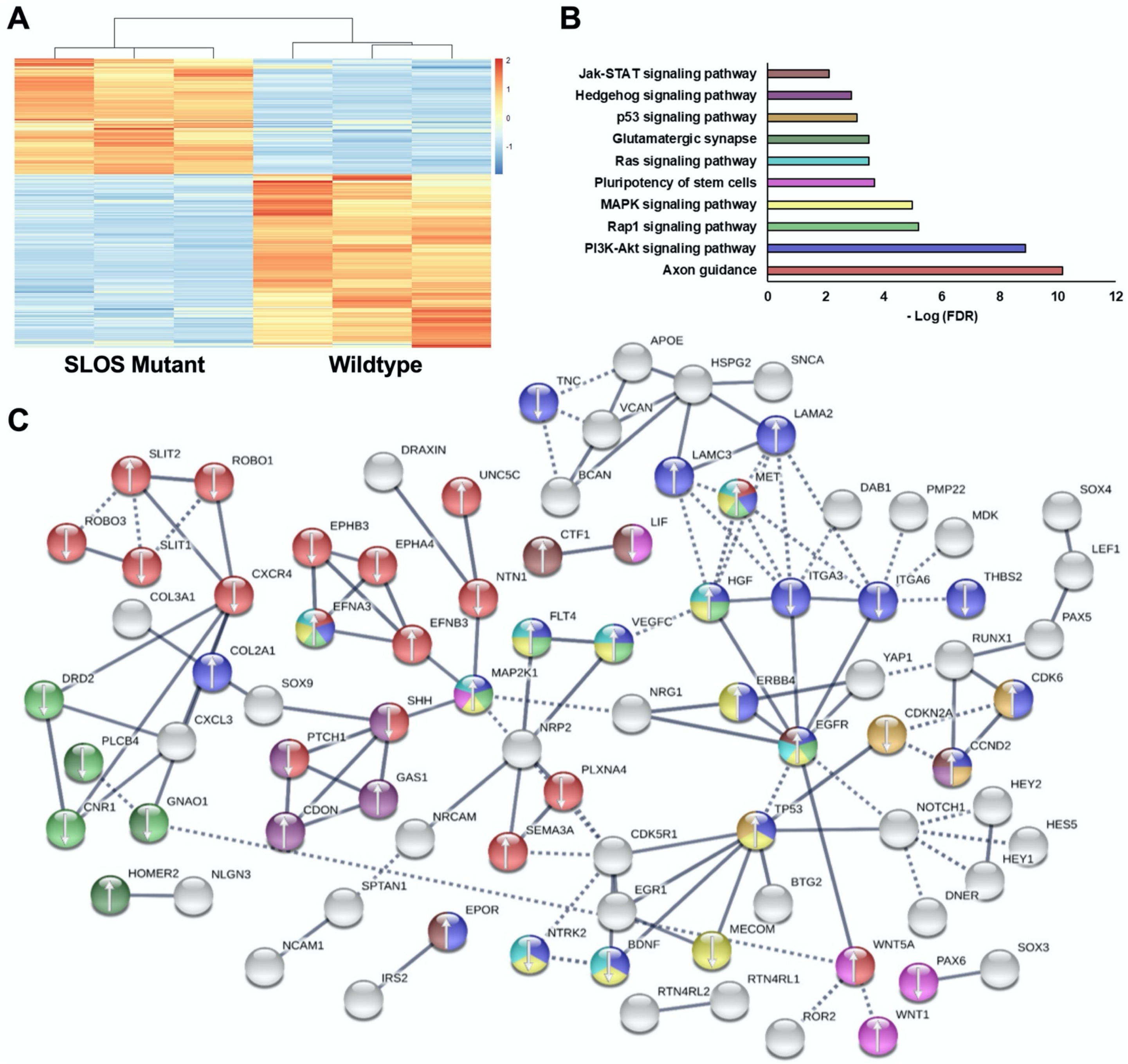


**Figure S1. Perturbation in *DHCR7* causes gene expression changes in neurogenic pathways in human NPCs.** (A) Hierarchical clustering heatmap of differentially expressed genes show distinct expression pattern changes in transcript abundance for SLOS mutant NPCs as compared to wild type. Red color represents increase in abundance, blue color represents relative decrease, and white color represents no change. (B) Enriched KEGG pathways identified among DEGs from SLOS mutant NPCs and involved in the Biological Function "development of the central nervous system", identified by Ingenuity Pathway Analysis (see Supporting Information). Color of bar corresponds to DEGs in the STRING network that are in enriched pathways. (C) String analysis of DEGs from SLOS mutant NPCs and involved in the Biological Function "Development of the central nervous system" identified by Ingenuity Pathway Analysis. Parameters of high confidence have been applied and only connected nodes are displayed. Arrows indicate whether the gene was up- or down-regulated. Color of DEG corresponds to enriched KEGG pathway. $n$ =3 biological replicates per genotype; DEGs met the criteria of fold-change > 1.5 between genotypes and adjusted $p$-value < 0.05.
